# *Drosophila* gustatory projections are segregated by taste modality and connectivity

**DOI:** 10.1101/2021.12.08.471796

**Authors:** Stefanie Engert, Gabriella R. Sterne, Davi D. Bock, Kristin Scott

## Abstract

Gustatory sensory neurons detect caloric and harmful compounds in potential food and convey this information to the brain to inform feeding decisions. To examine the signals that gustatory neurons transmit and receive, we reconstructed gustatory axons and their synaptic sites in the adult *Drosophila melanogaster* brain, utilizing a whole-brain electron microscopy volume. We reconstructed 87 gustatory projections from the proboscis labellum in the right hemisphere and 57 from the left, representing the majority of labellar gustatory axons. Gustatory neurons contain a nearly equal number of interspersed pre-and post-synaptic sites, with extensive synaptic connectivity among gustatory axons. Morphology- and connectivity-based clustering revealed six distinct groups, likely representing neurons recognizing different taste modalities. The vast majority of synaptic connections are between neurons of the same group. This study resolves the anatomy of labellar gustatory projections, reveals that gustatory projections are segregated based on taste modality, and uncovers synaptic connections that may alter the transmission of gustatory signals.

## Introduction

All animals have specialized sensory neurons dedicated to the detection of the rich variety of chemicals in the environment that indicate the presence of food sources, predators and conspecifics. Gustatory sensory neurons have evolved to detect food-associated chemicals and report the presence of caloric or potentially harmful compounds. Examining the activation and modulation of gustatory sensory neurons is essential as it places fundamental limits on the taste information that is funneled to the brain and integrated to form feeding decisions.

The *Drosophila melanogaster* gustatory system is an attractive model to examine the synaptic transmission of gustatory neurons. Molecular genetic approaches coupled with physiology and behavior have established five different classes of gustatory receptor neurons (GRNs) in adult *Drosophila* that detect different taste modalities. One class, expressing members of the Gustatory Receptor (GR) family including Gr5a and Gr64f, detects sugars and elicits acceptance behavior (Dahanukar et al 2001, Dahanukar et al 2007, Thorne et al 2004, Wang et al 2004). A second class expressing different GRs including Gr66a detects bitter compounds and mediates rejection behavior (Thorne et al 2004, Wang et al 2004, Weiss et al 2011). A third class contains the ion channel Ppk28 and detects water (Cameron et al 2010, Chen et al 2010). The fourth expresses the Ir94e ionotropic receptor and detects low salt concentrations, whereas the fifth contains the Ppk23 ion channel and may mediate detection of high salt concentrations (Jaeger et al 2018, Thistle et al 2012). In addition to well-characterized gustatory neurons and a peripheral strategy for taste detection akin to mammals (Yarmolinsky et al 2009), the reduced number of neurons in the *Drosophila* nervous system and the availability of electron microscopy (EM) brain volumes offer the opportunity to examine gustatory transmission with high resolution.

The cell bodies of gustatory neurons are housed in sensilla on the body surface including the proboscis labellum, an external mouthparts organ that detects taste compounds prior to ingestion (Stocker 1994). Gustatory neurons from each labellum half send bilaterally symmetric axonal projections to the subesophageal zone (SEZ) of the fly brain via the labial nerves. Gustatory axons terminate in the medial SEZ in a region called the anterior central sensory center (ACSC) (Hartenstein et al 2018, Miyazaki & Ito 2010, Thorne et al 2004, Wang et al 2004). Axons from bitter gustatory neurons send branches to the midline and form an interconnected medial ring whereas other gustatory axons remain ipsilateral and anterolateral to bitter projections. Although projections of different gustatory classes have been mapped using light level microscopy, the synaptic connectivity of gustatory axons in adult *Drosophila* is largely unexamined.

To explore the connectivity of GRNs and to lay the groundwork to study gustatory circuits with synaptic resolution, we used the recently available Full Adult Fly Brain (FAFB) Electron Microscopy (EM) dataset (Zheng et al 2018) to fully reconstruct gustatory axons and their synaptic sites. We reconstructed 87 GRN axonal projections in the right hemisphere and 57 in the left, representing between 83-96% and 54-63% of the total expected, respectively (Jaeger et al., 2018, Stocker, 1994). By annotating chemical synapses, we observed that GRNs contain a nearly equal number of interspersed pre-and post-synaptic sites. Interestingly, GRNs synapse onto and receive synaptic inputs from many other GRNs. Using morphology- and connectivity-based clustering, we identified six distinct neural groups, likely representing groups of GRNs that recognize different taste modalities. Our study reveals extensive anatomical connectivity between GRNs within a taste modality, arguing for pre-synaptic processing of taste information prior to transmission to downstream circuits.

## Results

### GRN axons contain pre-synaptic and post-synaptic sites

To systematically characterize gustatory inputs and outputs, we traced gustatory axons in the FAFB volume (Zheng et al 2018). Tracing was performed manually, using the annotation platform CATMAID (Saalfeld et al 2009). The GRNs from the proboscis labellum send axons through the labial nerve to the SEZ (Figure 1A). The labial nerve is a compound nerve, carrying sensory axons from the labellum, maxillary palp, and eye, as well as motor axons innervating proboscis musculature (Hampel et al 2020, Hartenstein et al 2018, Miyazaki & Ito 2010, Nayak & Singh 1983, Rajashekhar & Singh 1994). Different sensory afferents occupy different domains in the SEZ, with labellar gustatory axons terminating in the anterior central sensory center (ACSC) (Hartenstein et al 2018, Miyazaki & Ito 2010, Thorne et al 2004, Wang et al 2004). Therefore, to trace gustatory axons, we began by tracing neurites in the right labial nerve, readily identifiable in the EM dataset (Figure 1B and C), and selected fibers that terminated in the anterior central SEZ to trace to synaptic completion (Zheng et al 2018).

**Figure 1.**
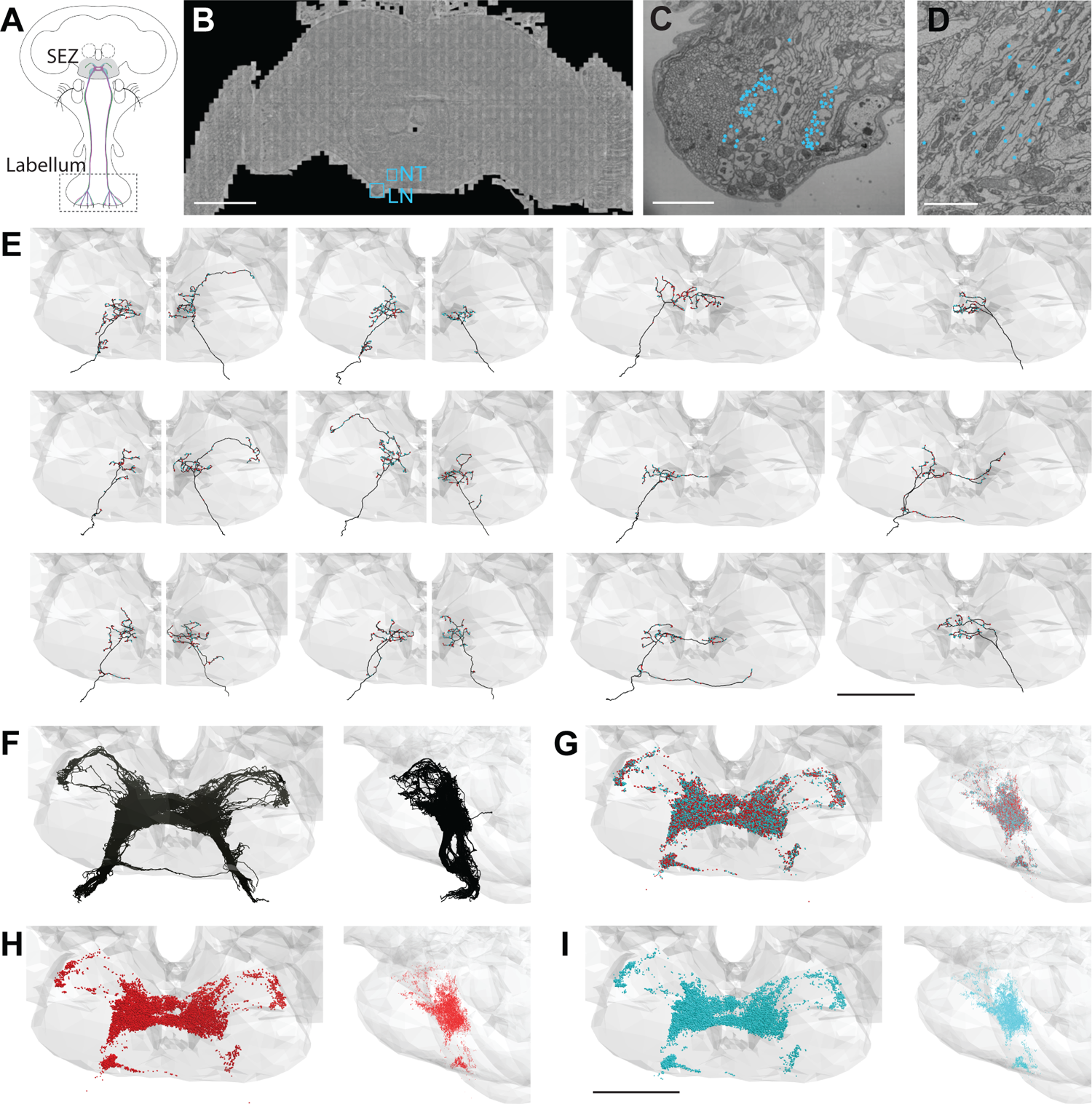
EM-based reconstructions of GRNs and synaptic sites. (A) Schematic showing GRNs in the proboscis labellum and their projections in the SEZ. (B) Location of the labial nerve (LN) and neural tract (NT) containing GRNs of the right hemisphere in the FAFB dataset (Z slice 3320, scale bar = 100 µM). (C) Cross-section of the labial nerve with traced GRNs indicated by asterisks (Z slice 3320, scale bar = 5 µM). (D) Neural tract with traced GRNs indicated by asterisks (Z slice 2770, scale bar = 5 µM). (E) Examples of reconstructed GRNs with presynaptic (red) and postsynaptic (blue) sites, scale bar = 50 µM. (F-I) Frontal and sagittal view of all reconstructed GRN axons (F), all presynaptic (red) and postsynaptic (blue) sites (G), presynaptic sites alone (H), and postsynaptic sites alone (I) Scale bar = 50 µM.

In tracing axons, we found that neurites with small to medium sized diameters in the dorsomedial labial nerve (Figure 1C) projected along a single neural tract (Figure 1D) to the anterior central region of the SEZ. This neural tract served as an additional site to select arbors for reconstruction. Individual fibers followed along the same tract and showed variation in terminal branching (Figure 1E). In total, we identified 87 axonal projections in the right hemisphere. Tracing from the left labial nerve and neural tract in the left hemisphere, we identified 57 additional projections. Misalignments in the EM volume precluded identification of additional GRNs in the left hemisphere. Because there are 90-104 GRNs per labellum (Jaeger et al 2018, Stocker 1994 we estimate that we have identified 83-96% of the GRN fibers from the right labellum and 54-63% from the left. The projections from the left and right labial nerves are symmetric and converge in a dense web in the anterior central SEZ (Figure 1F). This arborization pattern recapitulates the labellar sensory projections of the ACSC (Hartenstein et al 2018). We confirmed that the reconstructed neurites overlap with the known projection pattern of sugar and bitter GRNs in the registered fly brain template (Figure 1 - figure supplement 1) (Bogovic et al 2020), demonstrating that we have identified and traced GRNs.

In addition to the skeleton reconstructions, we manually annotated pre- and postsynaptic sites. The presence of T-shaped structures characteristic of presynaptic release sites (‘T bars’), synaptic vesicles, and a synaptic cleft were used to identify a synapse, consistent with previous studies (Zheng et al 2018). Synapses are sparse along the main neuronal tract and abundant at the terminal arborizations (Figure 1E). Each GRN has a large number of pre- and post-synaptic sites intermixed along the arbors (Figure 1E and G-I), characteristic of fly neurites (Bates et al 2020; Meinertzhagen 2018; Olsen & Wilson 2008; Takemura et al 2017). On average, a GRN contains 175 (± 6 SE) presynaptic sites and 168 ( ± 6 SE) postsynaptic sites, with individual GRNs showing wide variation in pre- and post-synapse number (Figure 1 - figure supplement 1B). GRNs are both pre- and post-synaptic to other GRNs, with each GRN receiving between 2% and 66% (average = 39%) of its synaptic input from other GRNs (Figure 1 - figure supplement 1C). The large number of synapses between GRNs suggests that communication between sensory neurons may directly regulate sensory output.

### Different GRN classes can be identified by morphology and connectivity

*Drosophila* GRNs comprise genetically defined, discrete populations that are specialized for the detection of specific taste modalities (Wang et al 2004, Cameron et al 2010, Jaeger et al 2018). As the EM dataset does not contain molecular markers to distinguish between GRNs recognizing different taste modalities, we set out to identify subpopulations of reconstructed GRNs based on their anatomy and connectivity.

We performed hierarchical clustering of GRN axons to define different subpopulations based on their morphology and synaptic connectivity. GRNs of the right hemisphere were used in this analysis as the dataset is more complete. Each traced skeleton was registered to a standard template brain (Bogovic et al 2018) and morphological similarity was compared pairwise using NBLAST in an all-by-all matrix (Costa et al 2016). Then, GRN-GRN connectivity was added for each GRN skeleton and the resulting merged matrix was min/max scaled. We then used Ward’s method to hierarchically cluster GRNs into groups (Ward 1963). We chose six groups as the number that minimizes within-cluster variance (Figure 2 - figure supplement 1A) (Braun et al 2010). Each group is composed of 7-23 GRNs that occupy discrete zones in the SEZ and share anatomically similar terminal branches (Figure 2).

**Figure 2.**
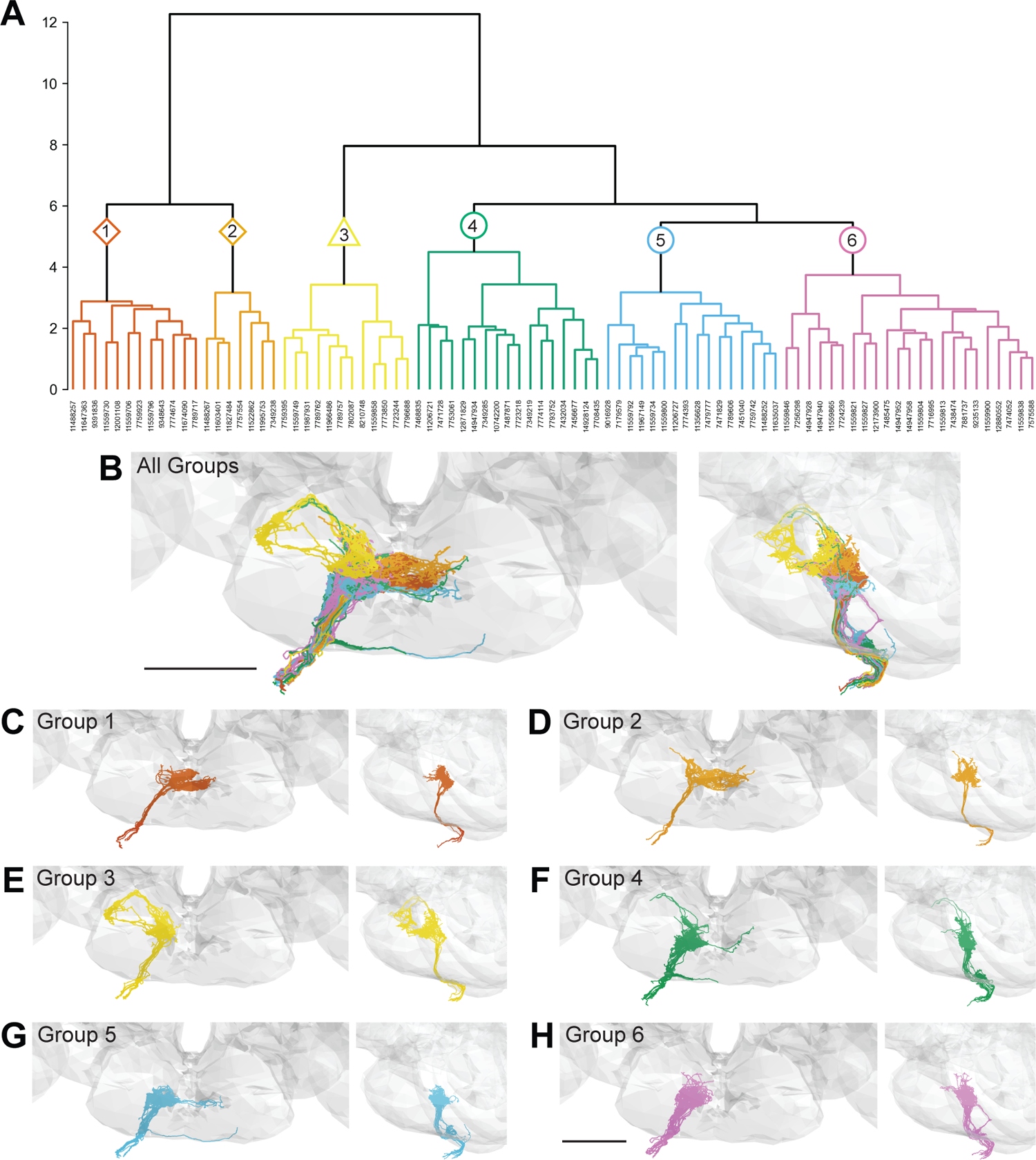
Morphology- and connectivity-based clustering generates distinct groups of GRNs. (A) Tree denoting relative similarity of GRNs based on morphology and connectivity of GRNs in the right hemisphere. (B) Frontal and sagittal view of all GRN groups, colored according to A. (C-H) Frontal and sagittal view of group 1 - group 6 GRNs, scale bar = 50 µM.

To evaluate whether the different groups represent GRNs detecting different taste modalities, we compared the anatomy of each group in the right hemisphere with that of known GRN classes, using NBLAST similarity scores. We registered EM reconstructed GRN projections and GRN projections from immunostained brains to the same standard brain template for direct comparisons (Bogovic et al 2020). For each group, we performed pairwise comparisons against bitter (Gr66a; Wang et al 2004, Thorne et al 2004), sugar (Gr64f; Dahanukar et al 2007), water (Ppk28; Cameron et al 2010, Chen et al 2010), and low salt (Ir94e; Croset et al 2016, Jaeger et al 2018) projections. There is not a specific genetic marker for high salt projections, as Ppk23 labels both bitter and high salt GRNs (Jaeger et al 2018). These comparisons (Methods, NBLAST analysis for taste modality assignment) revealed that group 1 and group 2 best match bitter projections, forming a characteristic medial ringed web. Group 3 projections show greatest similarity to low salt GRNs, with distinctive dorsolateral branches. Groups 4, 5 and 6 are anatomically very similar, and identity assignments are tentative. Group 6 best matches water GRNs. Group 4 and group 5 best match sugar GRNs. Because group 4 shows greater similarity with sugar GRNs based on NBLAST scores and because it contains a dorsolateral branch seen in Gr64f projections and not seen in group 5 projections, we hypothesize that group 4 is composed of sugar GRNs and that the remaining group 5 is composed high salt GRNs. Comparison of each group with its GRN category best match in the 3-dimensional standard fly brain template supports the view that Group 1 and 2 are bitter GRNs, group 3 low-salt, group 4 sugar, and group 6 water GRNs (Figure 3). Thus, morphological and connectivity clustering suggests molecular and functional identities of different GRNs.

**Figure 3.**
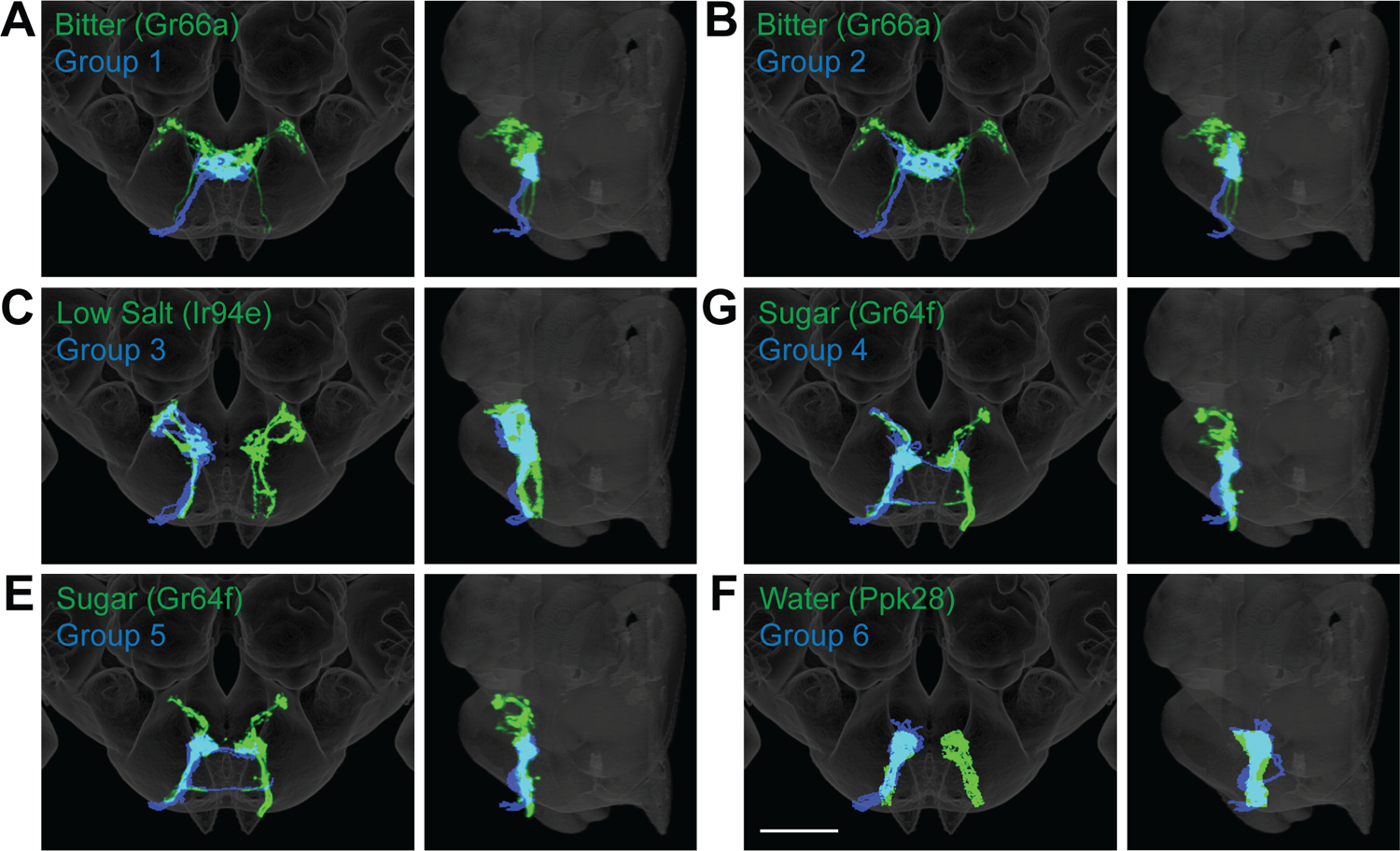
Anatomy of different GRN groups overlays with GRNs of different taste categories. NBLAST comparisons yielded best matches of EM groups and GRNs of different taste classes. A-F. Overlain are EM Groups 1-6 (blue) and best NBLAST match (green), frontal view (left) and sagittal view (right), scale bar = 50 µM.

An identical clustering analysis of GRNs from the left hemisphere yielded 7 groups of 4-15 neurons (Figure 3 – figure supplement 1-2). Groups 1 and 2 best match bitter projections and group 6 best matches low salt projections (Methods, NBLAST analysis for taste modality assignment), with anatomy consistent with known projection patterns. Other groups are not well-resolved (Methods, NBLAST analysis for taste modality assignment), arguing that a more complete dataset is necessary to resolve GRN categories in the left hemisphere.

### GRNs are highly interconnected via chemical synapses

As GRNs have a large number of synaptic connections with other GRNs (Figure 1 - figure supplement C), we examined whether synapses exist exclusively between neurons of the same group, likely representing the same taste modality, or between multiple groups. The all-by-all connectivity matrix illustrated blocks of connectivity within groups, with fewer connections between groups (Figure 4A). To quantify this, we summed all GRN-GRN connections within and between groups. This analysis revealed that most synapses are between neurons of the same group (79%), while only 21% of the synapses are between GRNs of different groups (Figure 4B). For example, group 4 neurons receive 1468 synapses from other group 4 neurons and 38 from group 3, 156 from group 5, and 130 from group 6 neurons. Focusing on connections of five or more synapses between GRN pairs, representing high confidence connections (Buhmann et al 2021, Li et al 2020a, Takemura et al 2013, Takemura et al 2015), resulted in the elimination of some but not all between-group connections (Figure 4C), with between-group connections representing only 10% of all GRN connections.

**Figure 4.**
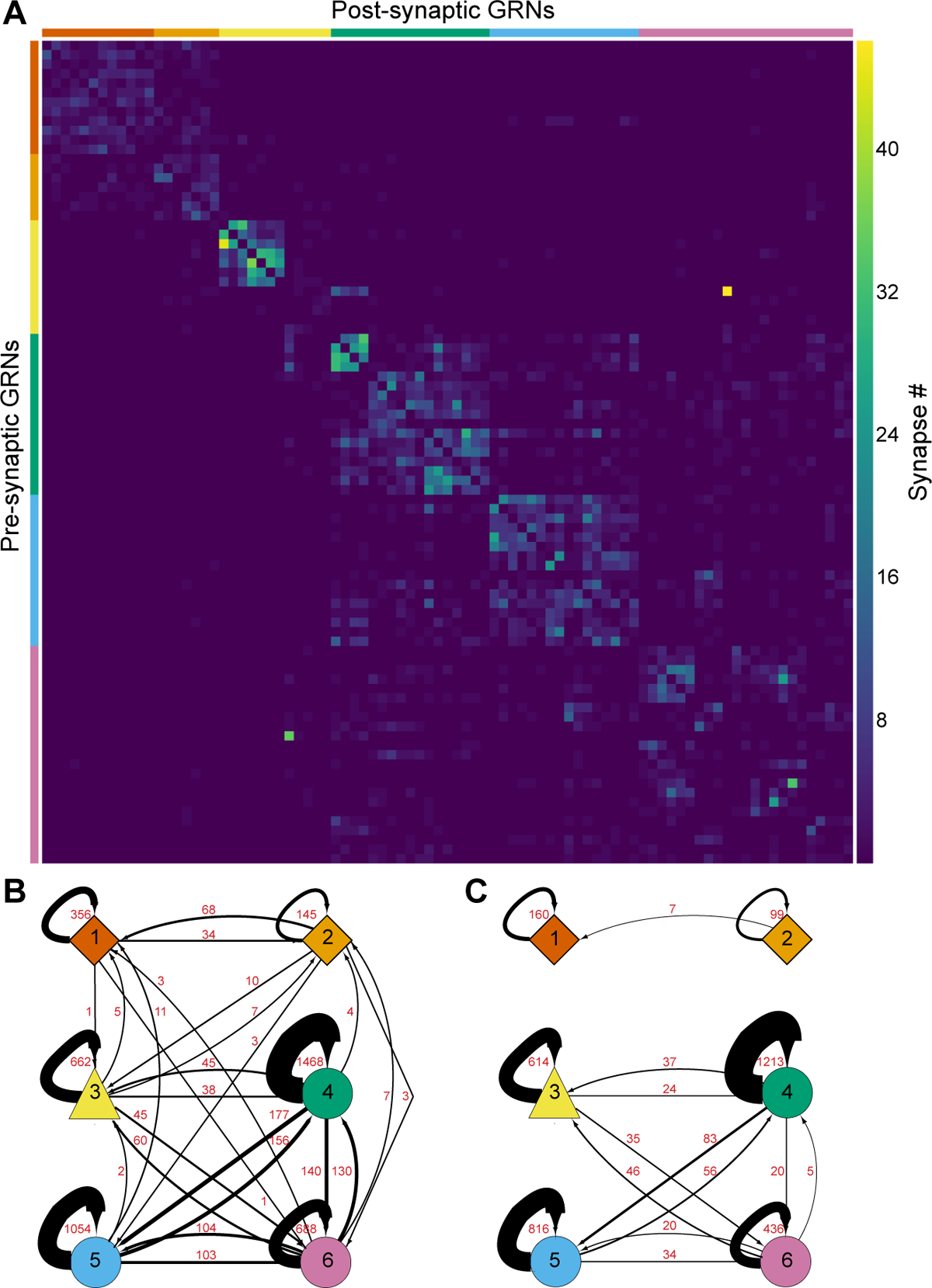
GRNs are highly interconnected via chemical synapses. (A) Connectivity matrix of GRNs in the right hemisphere. GRNs groups are color-coded and ordered according to Figure 2. Color coding within the matrix indicates the number of synapses from the pre- to the post-synaptic neuron, indicated in the legend. (B) Connectivity between GRN groups. Colors correspond to groups in Figure 2. Arrow thickness scales with the number of synapses, indicated in red. (C) Connectivity between GRN groups as in B, showing only connections of 5 or more synapses.

The large numbers of chemical synapses between GRNs within a group may provide a mechanism to amplify signals of the same taste modality. In contrast, weak connectivity between GRNs of different groups may serve to integrate taste information from different modalities before transmission to downstream circuitry. We note that misclassification of individual GRNs in the clustering analysis may result in over- or underestimates of GRN connectivity within and between groups.

### Interactions between sugar and water GRNs are not observed by calcium or voltage imaging

To examine whether the small number of connections between GRNs of different taste modalities results in cross-activation of GRNs detecting different primary tastant classes, we tested if activation of one GRN class results in propagation of activity to other GRN classes *in vivo*. As the connectivity data suggests that sugar and water GRNs are weakly connected (Figure 4 B-C, group 4 and group 6), we wondered if appetitive GRNs might be interconnected to amplify appetitive signals to downstream feeding circuits. To test for interactions between appetitive GRNs, we undertook calcium and voltage imaging studies in which we monitored the response of a GRN class upon activation of other GRN classes.

We expressed the calcium indicator GCaMP6s in genetically defined sugar, water or bitter sensitive GRNs to monitor excitatory responses upon artificial activation of different GRN classes. To ensure robust and specific activation of GRNs, we expressed the mammalian ATP receptor P2X2 in sugar, water or bitter GRNs, and activated the GRNs with an ATP solution presented to the fly proboscis while imaging gustatory projections in the brain (Yao et al 2012, Harris et al 2015). Expressing both P2X2 and GCaMP6s in sugar, water or bitter GRNs elicited strong excitation upon ATP presentation (Figure 5A-B and G-H and Figure 5 - figure supplement 1-3 C-D), demonstrating the effectiveness of this method. As bitter cells are synaptically connected to each other but not to sugar or water cells, we hypothesized that they would not be activated by sugar or water GRN activation. Consistent with the EM connectivity, activation of sugar or water GRNs did not activate bitter cells, nor did bitter cell activation elicit responses in sugar or water axons (Figure 5 - figure supplement 1E-H; Figure 5 - figure supplement 2E-F; Figure 5 - figure supplement 3G-H). In contrast, the EM connectivity indicates possible interactions between sugar and water GRNs. However, we did not observe responses in sugar GRNs upon water GRN activation (Figure 5C-D; Figure 5 - figure supplement 2IJ) or responses in water GRNs upon sugar GRN activation (Figure 5I and J; Figure 4 - figure supplement 3E and F). To examine whether interactions between modalities are modulated by the feeding state of the fly, we performed the activation and imaging experiments in both fed and starved flies (Figure 5 - figure supplement 1-6). These experiments did not reveal feeding state-dependent interactions between GRN populations.

**Figure 5.**
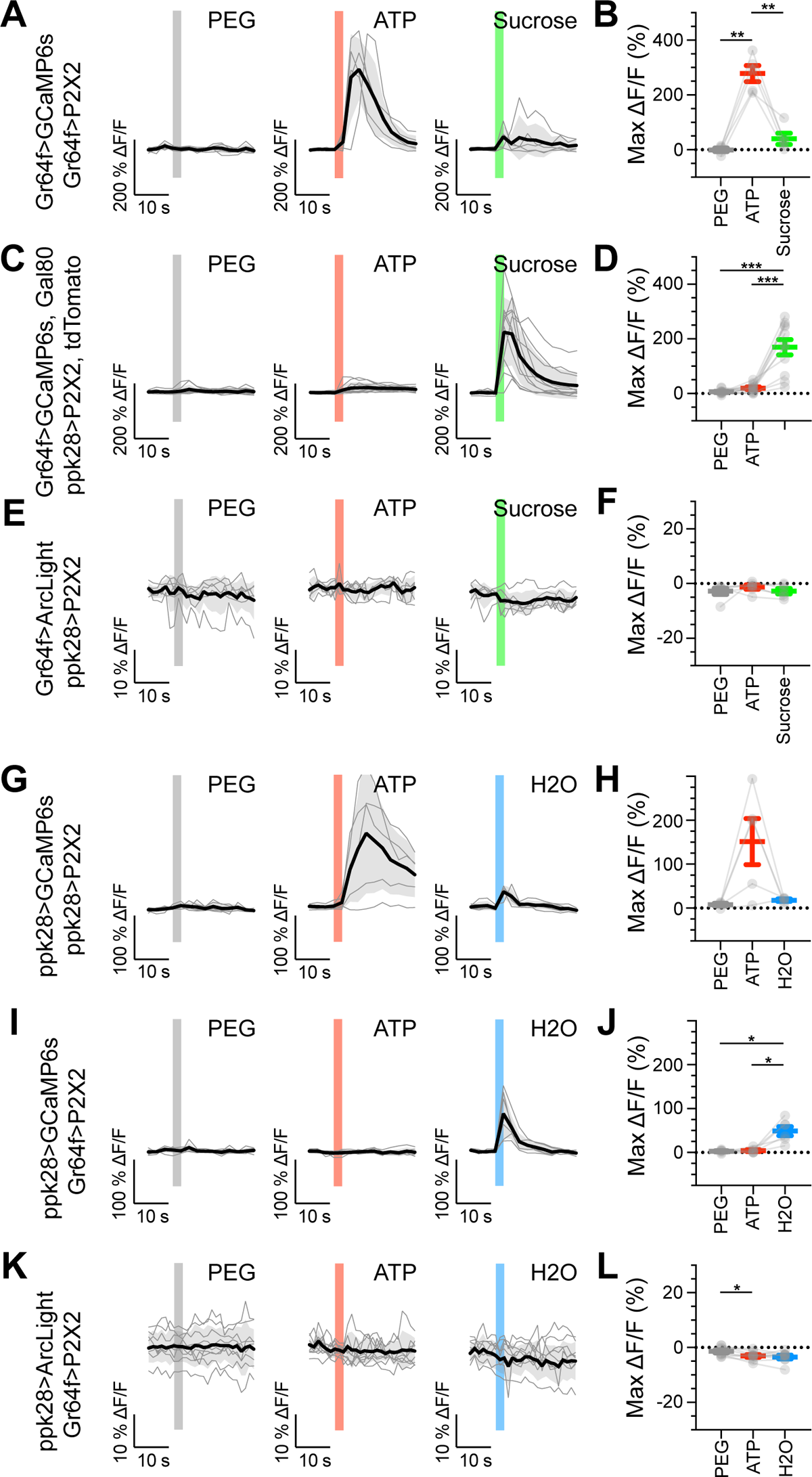
Sugar and water GRNs do not activate each other. (A, B) Calcium responses of sugar GRNs expressing P2X2 and GCaMP6s to proboscis presentation of PEG as a negative control, ATP to activate P2X2, or sucrose as a positive control. GCaMP6s fluorescence traces (ΔF/F) (A) and maximum ΔF/F post stimulus presentation (B), n = 5. Sugar GRNs responded to ATP, but the response to subsequent sucrose presentation was attenuated. (C, D) GCaMP6s responses of sugar GRNs in flies expressing P2X2 in water GRNs to PEG, ATP, and sucrose delivery, ΔF/F traces (C) and maximum ΔF/F graph (D), n = 11. (E, F) ArcLight responses of sugar GRNs in flies expressing P2X2 in water GRNs, ΔF/F traces (E) and maximum ΔF/F graph (F), n = 6. (G, H) Calcium responses of water GRNs expressing P2X2 and GCaMP6s to proboscis delivery of PEG (negative control), ATP, and water (positive control), ΔF/F traces (G) and maximum ΔF/F graph (H), n = 5. Water GRNs responded to ATP presentation, but the subsequent response to water was diminished. (I, J) GCaMP6s responses of water GRNs in flies expressing P2X2 in sugar GRNs to PEG, ATP, and water, ΔF/F traces (I) and maximum ΔF/F graph (J), n = 6. (K, L) ArcLight responses of water GRNs in flies expressing P2X2 in sugar GRNs to PEG, ATP, and water, ΔF/F traces (K) and maximum ΔF/F graph (L), n = 9. For all traces, stimulus presentation is indicated by shaded bars. Traces of individual flies to the first of three taste stimulations (shown in Figure 5 – Figure supplements 2, 3 and 7) are shown in grey, the average in black, with the SEM indicated by the grey shaded area. Repeated measures ANOVA with Tukey’s multiple comparisons test, *p<0.05, **p<0.01, ***p<0.001.

We reasoned that interactions between sugar and water GRNs might be inhibitory, providing a mechanism to weight different appetitive taste inputs. To examine this, we expressed the voltage indicator ArcLight (Cao et al 2013), which reliably reports hyperpolarization, in sugar GRNs while activating water GRNs via P2X2 and vice versa. These experiments revealed no change in voltage in one appetitive gustatory class upon activation of the other (Figure 5E-F and K-L: Figure 4 - figure supplement 7). Overall, despite the potential for crosstalk between different modalities revealed by EM, we observed no communication between appetitive GRNs by calcium or voltage imaging of gustatory axons.

## Discussion

In this study, we characterized different classes of gustatory projections and their interconnectivity by high-resolution EM reconstruction. We identified different projection patterns corresponding to gustatory neurons recognizing different taste modalities. The extensive connections between GRNs of the same taste modality provide anatomical evidence of pre-synaptic processing of gustatory information.

An emerging theme stemming from EM reconstructions of *Drosophila* sensory systems is that sensory neurons of the same subclass are synaptically connected. In general, different sensory neuron subclasses have spatially segregated axonal termini in the brain, thereby constraining the potential for connectivity. In the adult olfactory system, approximately 40% of the input onto olfactory receptor neurons (ORNs) comes from other ORNs projecting to the same olfactory glomerulus (Horne et al 2018, Schlegel et al 2021, Tobin et al 2017). Similarly, mechanosensory projections from the Johnston’s Organ of the same submodality are anatomically segregated and synaptically connected (Hampel et al 2020). In *Drosophila* larvae, 25% of gustatory neuron inputs are from other GRNs, although functional classes were not resolved (Miroschnikow et al 2018). In the adult *Drosophila* gustatory system, we also find that GRNs are interconnected, with approximately 39% of GRN input coming from other GRNs. Consistent with other classes of sensory projections, we find that gustatory projections are largely segregated based on taste modality and form connected groups. A general function of sensory-sensory connections seen across sensory modalities may be to enhance weak signals or to increase dynamic range.

By clustering neurons based on anatomy and connectivity, we were able to resolve different GRN categories. The distinct morphologies of bitter neurons and low salt-sensing neurons, known from immunohistochemistry, are recapitulated in the projection patterns of GRN groups 1, 2 and 3 of the right hemisphere, enabling high-confidence identification. It is interesting that bitter projections cluster into two distinct groups, suggesting different subsets. We hypothesize that these reflect bitter GRNs from different taste bristle classes or bitter GRNs with different response properties (Dweck and Carlson 2020). The projections of high salt, sugar and water-sensing neurons are ipsilateral, with similarities in their terminal arborizations (Jaeger et al 2018, Wang et al 2004). Nevertheless, comparisons between EM and light-level projections argue that these taste categories are also resolved into different, identifiable clusters. However, as these categories are based on anatomical comparisons alone, they remain tentative until further examination of taste response profiles of connected second-order neurons, may now be identified by EM tracing downstream of the reconstructed GRNs reported here.

Examining GRN-GRN connectivity revealed connectivity between GRNs of the same group as well as different groups. While it is tempting to speculate that interactions between different taste modalities may amplify or filter activation of feeding circuits, we were unable to identify cross-activation between sugar and water GRNs by calcium or voltage imaging. It is possible that these interactions are dependent on a feeding state or act on a timeframe not examined in this study. Alternatively, activation may fall below the detection threshold of calcium or voltage imaging. Additionally, far fewer synapses occur between anatomical classes than within classes, especially restricting analyses to neurons connected by 5 or more synapses (Figure 4C), suggesting that the small number of synapses may not be relevant for taste processing. Finally, the anatomy and connectivity-based clustering may not categorize all individual GRNs correctly, and misclassification of GRNs would impact connectivity analyses. Regardless, our studies suggest that pre-synaptic connectivity between different GRN classes does not substantially contribute to taste processing.

Overall, this study resolves the majority of labellar gustatory projections and their synaptic connections, revealing that gustatory projections are segregated based on taste modality and synaptic connections. The identification of GRNs detecting different taste modalities now provides an inroad to enable the examination of the downstream circuits that integrate taste information and guide feeding decisions.

## Materials and Methods

### Key Resources Table

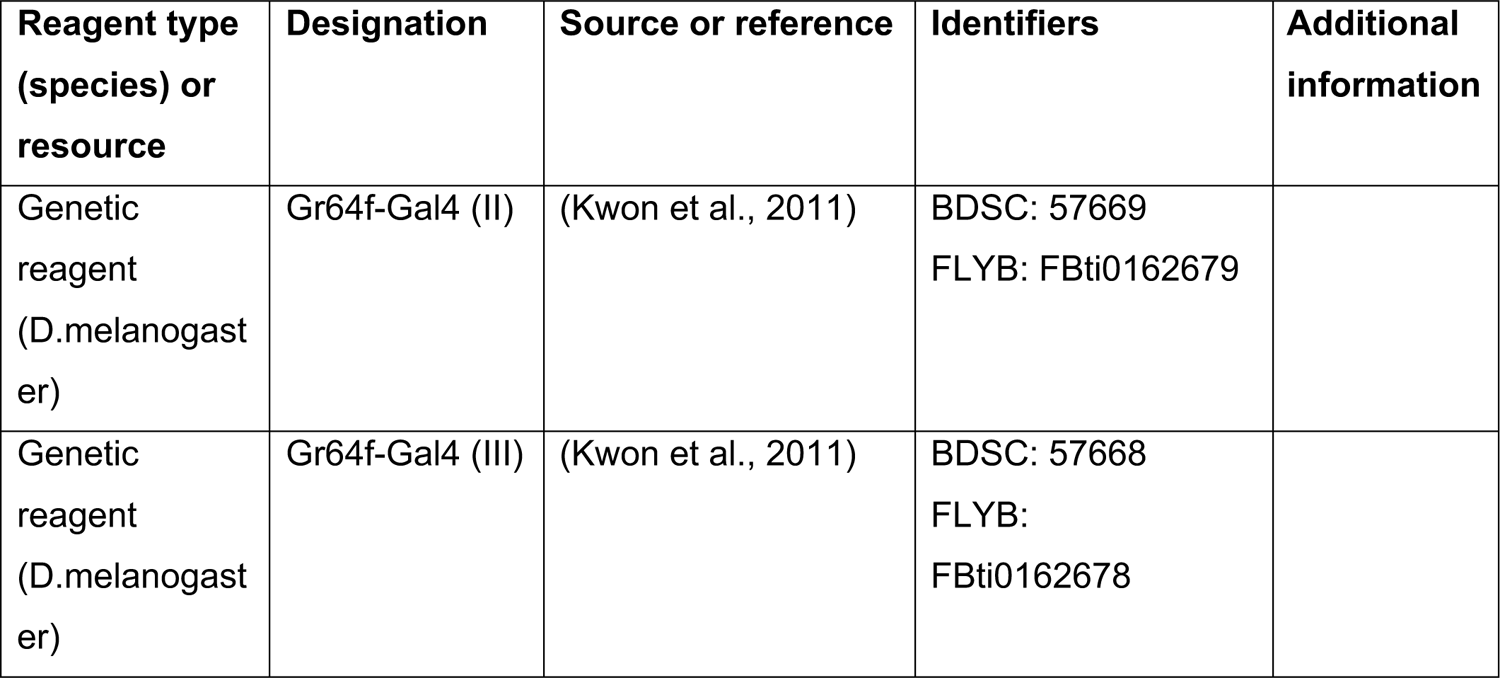
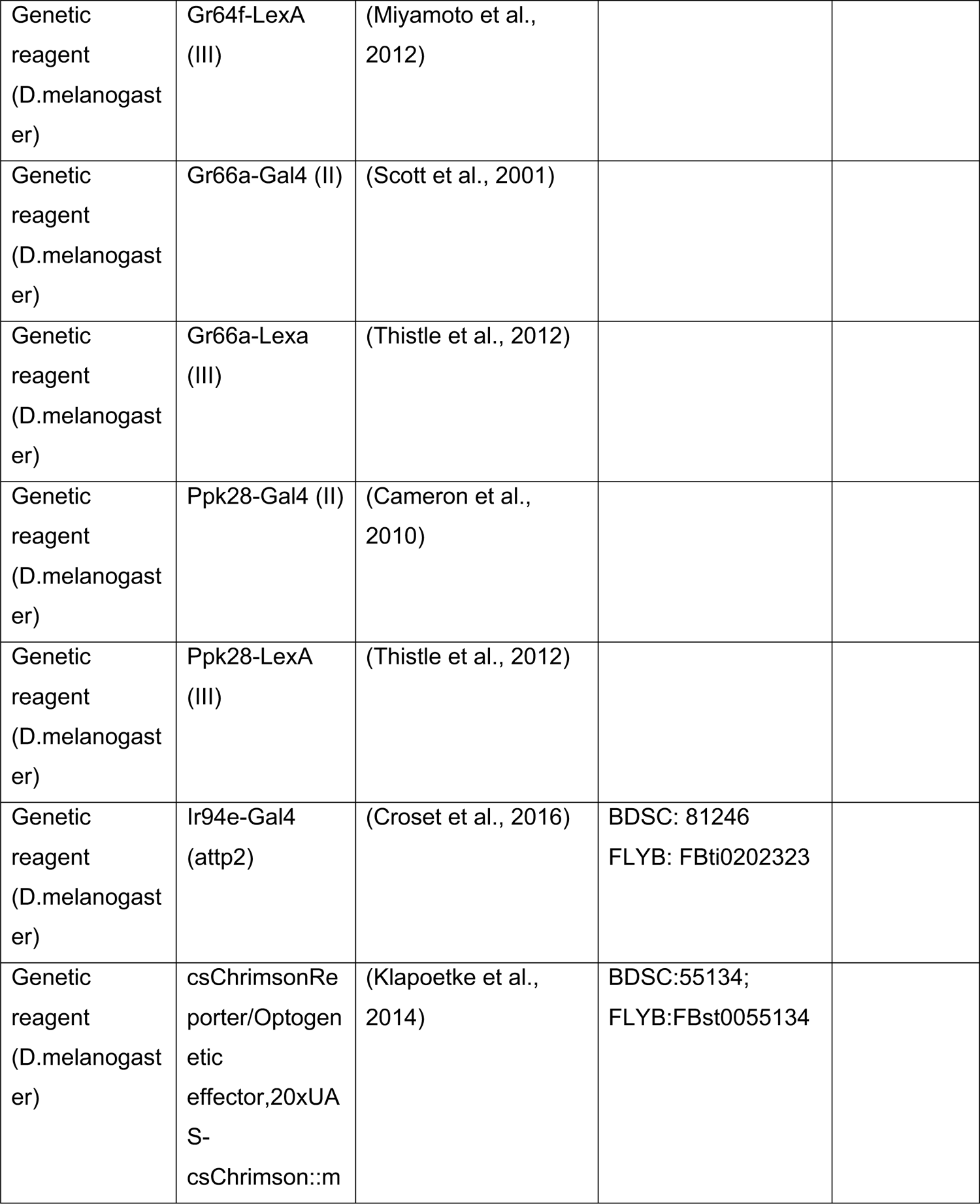
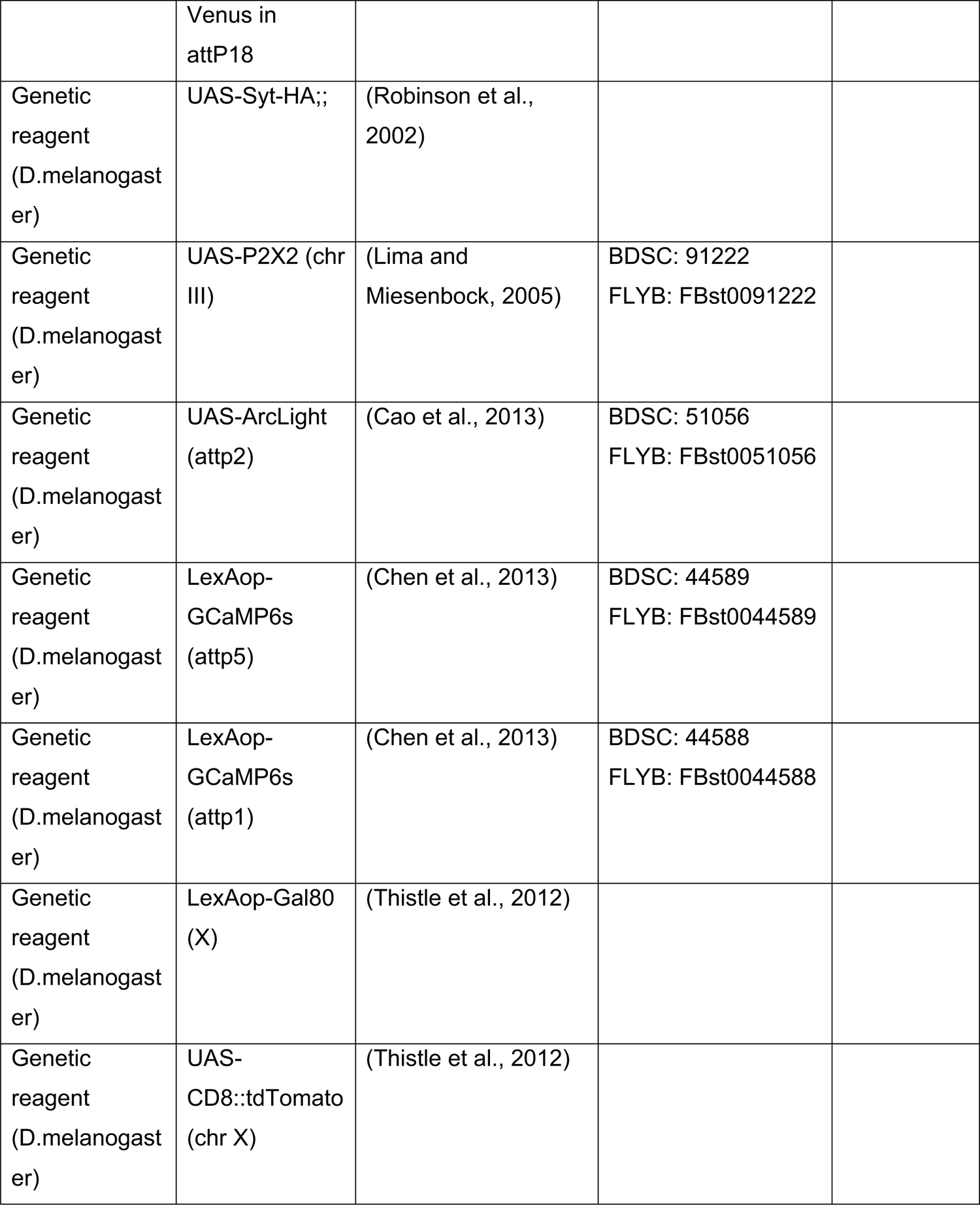
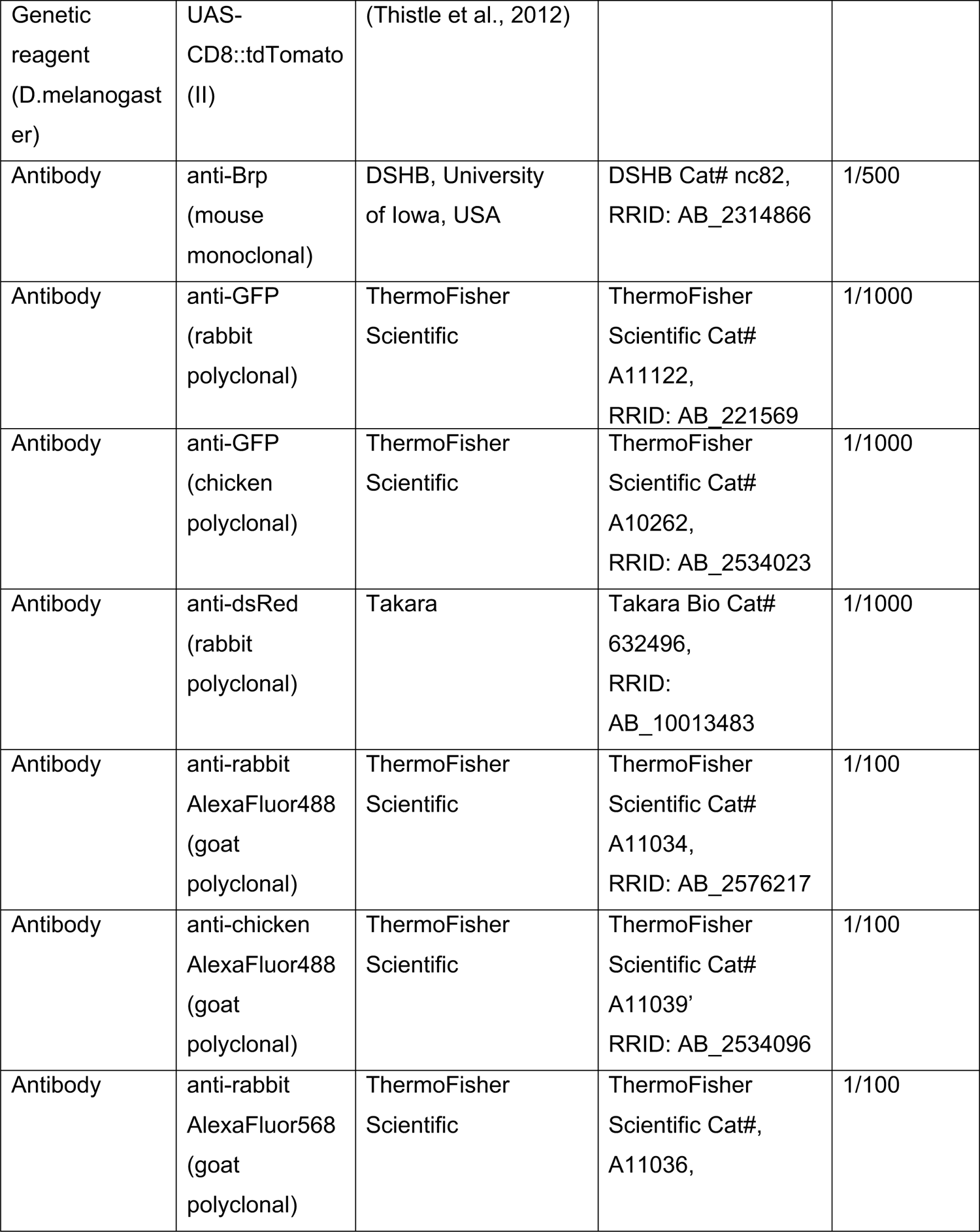
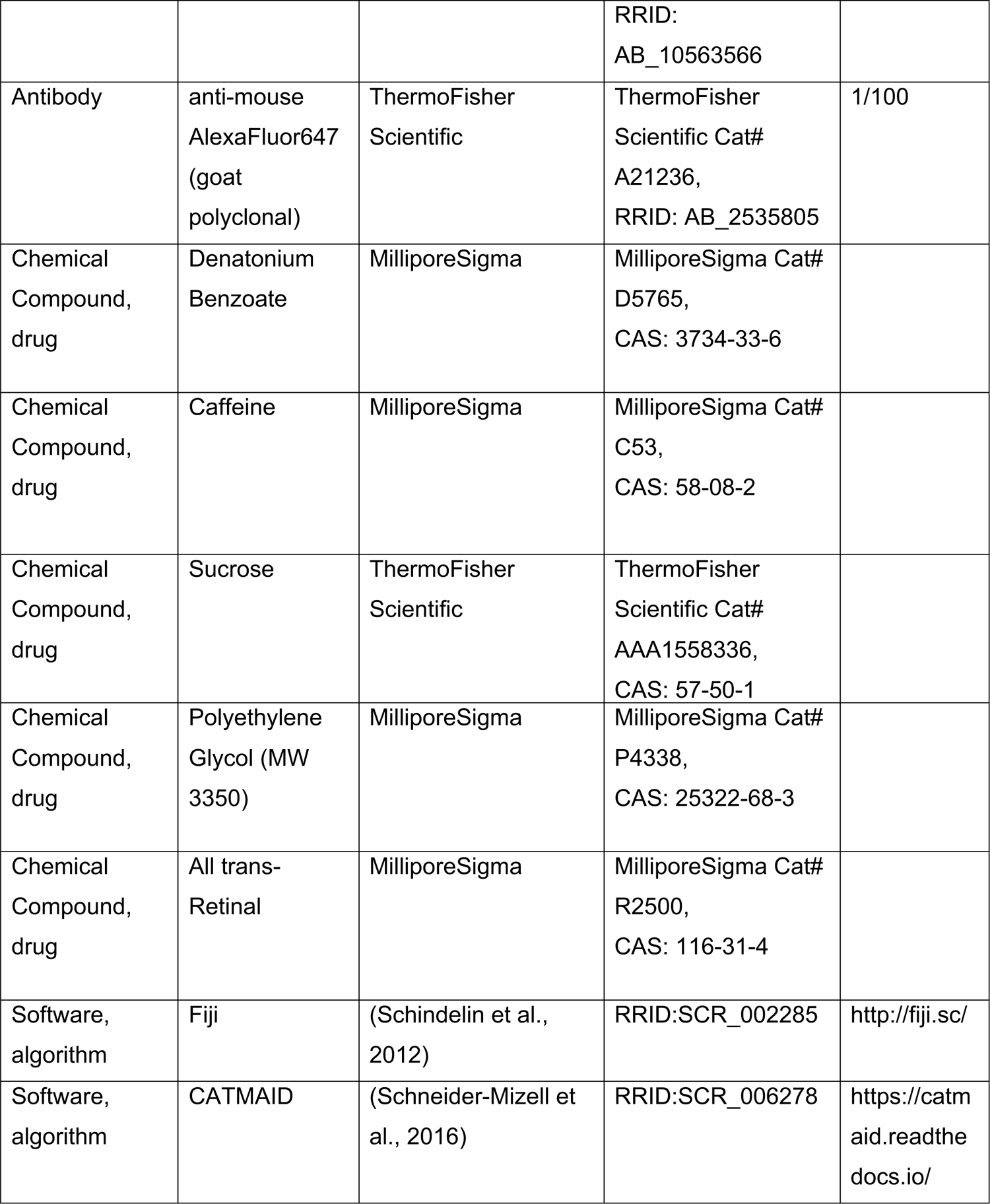
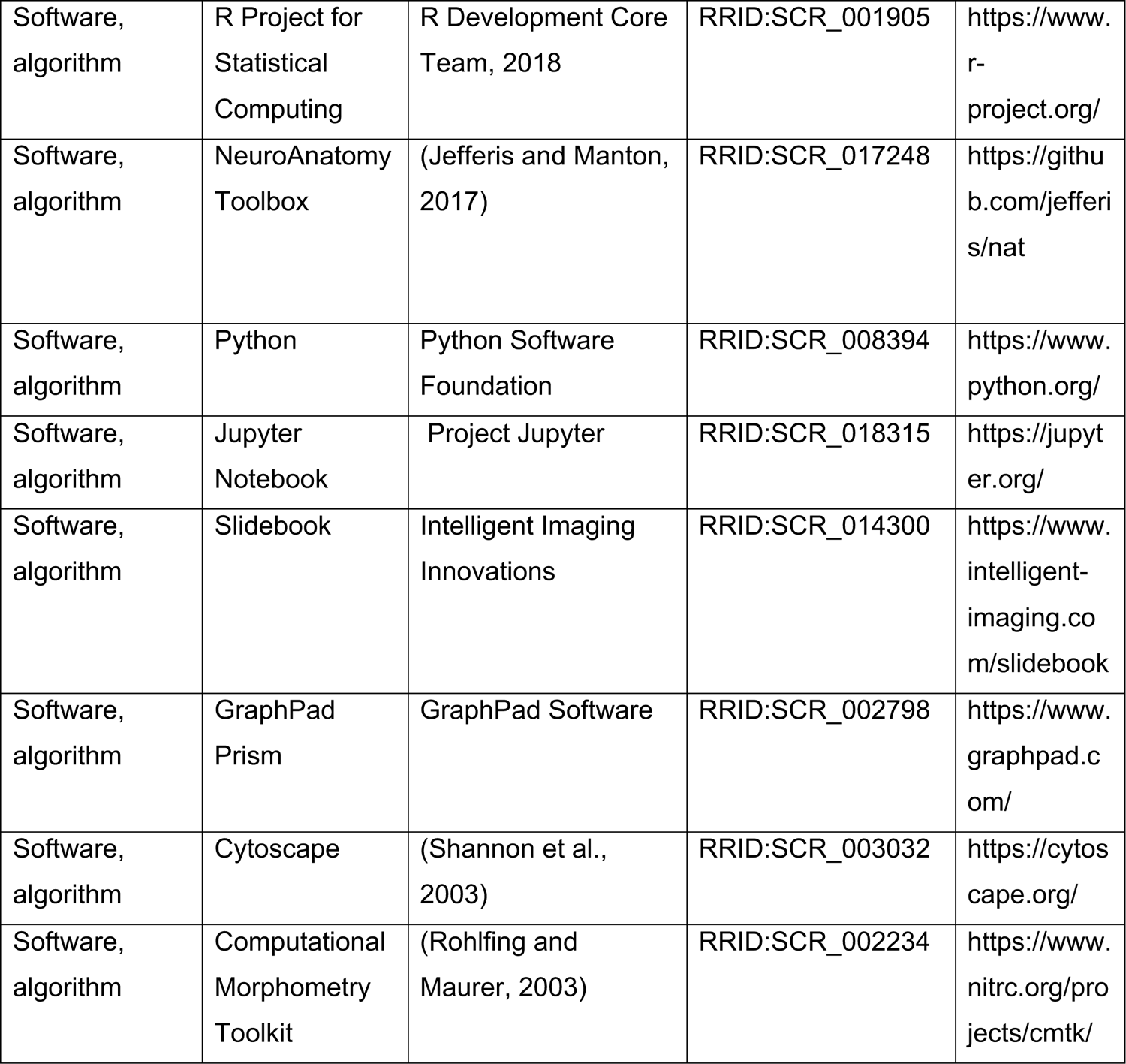

### Experimental Animals

Experimental animals were maintained on standard agar/molasses/cornmeal medium at 25°C. For imaging experiments requiring food-deprived animals, flies were placed in vials containing wet kimwipes for 23-26 hours prior to the experiment. For behavioral experiments, flies were placed on food supplemented with 400μM trans-retinal for 24 hours prior to the experiment.

### EM reconstruction

Neuron skeletons were reconstructed in a serial sectioned transmission electron microscopy dataset of the whole fly brain (Zheng et al 2018) using the annotation software CATMAID (Saalfeld et al 2009). GRN projections were identified based on their extension into the labial nerve and localization to characteristic neural tracts in the SEZ. Skeletons were traced to completion either entirely manually or using a combination of an automated segmentation (Li et al 2020b) and manual tracing as previously described (Hampel et al 2020). Chemical synapses were annotated manually and neurons were traced to synaptic completion, using criteria previously described (Zheng et al 2018). Skeletons were reviewed by a second specialist, so that the final reconstruction presents the consensus assessment of at least two specialists. Skeletons were exported from CATMAID as swc files for further analysis, and images of skeletons were exported directly from CATMAID. FAFB neuronal reconstructions will be available from Virtual Fly Brain (https://fafb.catmaid.virtualflybrain.org/).

### Clustering of GRNs

GRNs were hierarchically clustered based on morphology and connectivity using NBLAST and synapse counts. First, GRN skeletons traced in FAFB were registered to the JRC2018U template (Bogovic et al 2018) and compared in an all-by-all fashion with NBLAST (Costa et al. 2016). NBLAST analysis was carried out with the natverse toolkit in R (Bates et al. 2020; R Development Core Team, https://www.r-project.org/). The resulting matrix of NBLAST scores was merged with a second matrix containing all-by-all synaptic connectivity counts for the same GRNs. The resulting merged matrix was min-max normalized such that all values fall within the range of 0 and 1. The merged, normalized matrix was hierarchically clustered using Ward’s method (Ward 1963) in Python (Python Software Foundation, https://www.python.org/) with SciPy (Virtanen et al 2020). The number of groups was chosen based on analysis of Ward’s joining cost and the differential of Ward’s joining cost. Connectivity data of GRNs was exported from CATMAID for further analysis and connectivity diagrams were generated using CytoScape (Shannon et al 2003).

### NBLAST analysis for taste modality assignment

GRN skeletons traced in FAFB were registered to the JRC2018U template and summed in FIJI to create a composite stack of the combined morphologies of all individual GRNs in a given group (as assigned by morphology and connectivity clustering). The morphology of the composite stack for each group was compared to an image library of GRN projection patterns using NBLAST (Costa et al. 2016). The image library contained projection patterns of Gr66a-GAL4, Gr64f-GAL4, Ir94e-GAL4, and Gr64f-GAL4 brains, 3 per genotype, registered to the JRC2018U template, prepared as described (see the “Immunohistochemistry” section below). Group identity was assigned based on the top hit from the image library. Following NBLAST analysis, the anatomy of each group was compared to the projection pattern of its top hit using VVDViewer.

NBLAST of groups in the right hemisphere against known GRN categories yielded the following top GRN matches, (NBLAST score): Group 1, Gr66a-GAL4 #1 (47367); Group 2, Gr66a-GAL4 #1 (55586); Group 3, Ir94e-GAL4 #2 (67719); Group 4, Gr64f-GAL4 #2 (65797); Group 5, Gr64f-GAL4 #2 (56161); Group 6, Ppk28-GAL4 #1 (58018). NBLAST of groups in the left hemisphere against known GRN categories yielded the following top GRN matches, (NBLAST score): Group 1, Gr66a-GAL4 #3 (36848); Group 2, Gr66a-GAL4 #3 (34344); Group 3, Gr64f-GAL4 #2 (10776); Group 4, Gr64f-GAL4 #2 (43049); Group 5, Gr64f-GAL4 #2 (18544); Group 6, Ir93a-GAL4 #2 (22987).; Group 7, Gr66a-GAL4 #2 (48780).

### Calcium and Voltage Imaging Preparation

For imaging studies of GRNs, mated females, 10 to 21 days post eclosion, were dissected as previously described (Harris et al 2015), so that the brain was submerged in artificial hemolymph (AHL) (Wang et al 2003) while the proboscis was kept dry and accessible for taste stimulation. To avoid occlusion of taste projections in the SEZ, the esophagus was cut. The front legs were removed for tastant delivery to the proboscis. AHL osmolality was assessed as previously described (Jourjine et al 2016) and adjusted according to the feeding status of the animal. In fed flies, AHL of ∼250mOsm was used (Wang et al 2003). The AHL used for starved flies was diluted until the osmolality was ∼180mOsm, consistent with measurements of the hemolymph osmolality in food deprived flies (Jourjine et al 2016).

### Calcium Imaging

Calcium transients reported by GCaMP6s and GCaMP7s were imaged on a 3i spinning disk confocal microscope with a piezo drive and a 20x water immersion objective (NA=1). For our studies of GRNs, stacks of 14 z-sections, spaced 1.5 microns apart, were captured with a 488nm laser for 45 consecutive timepoints with an imaging speed of ∼0.3 Hz and an optical zoom of 2.0. For better signal detection, signals were binned 8×8, except for Gr64f projections, which underwent 4×4 binning.

### Voltage Imaging

Voltage responses reported by ArcLight were imaged similarly to the calcium imaging studies. To increase imaging speed, the number of z planes was reduced to 10, and the exposure time was decreased from 100ms to 75ms, resulting in an imaging speed of ∼0.7Hz. To maintain a time course comparable to that of the calcium imaging experiments of GRNs, the number of timepoints was increased to 90. Signals were binned 8×8 in each experiment.

### Taste stimulations

Taste stimuli were delivered to the proboscis via a glass capillary as previously described (Harris et al 2015). For GRN studies, each fly was subjected to three consecutive imaging sessions, each consisting of a taste stimulation at time point 15, 25 and 35 (corresponding to 30, 50.5, 71.5 sec). During the first imaging session, the fly was presented with a tasteless 20% polyethylene glycol (PEG, average molecular weight 3350 g/mol) solution, acting as a negative control. PEG was used in all solutions except water solutions, as this PEG concentration inhibits activation of water GRNs (Cameron et al 2010). This was followed in the second session with stimulations with 100mM ATP in 20%PEG. In the last imaging session, each fly was presented with a tastant acting as a positive control in 20% PEG (Gr64f: 1M sucrose; Gr66a: 100mM caffeine, 10mM denatonium benzoate; ppk28: H2O; ppk23: 1M KCl in 20% PEG).

### Imaging Analysis

Image analysis was performed in FIJI (Schindelin et al 2012). Z stacks for each time point were converted into maximum z-projections for further analysis. After combining these images into an image stack, they were aligned using the StackReg plugin in FIJI to correct for movement in the xy plane (Thevenaz et al 1998).

For our exploration of interactions between GRN subtypes, one ROI was selected encompassing the central arborization of the taste projection in the left or right hemisphere of the SEZ in each fly. Whether the projection in the left or right hemisphere was chosen depended on the strength of their visually gauged response to the positive control. The exception was Gr66a projections, in which the entire central projection served as ROI. If projections did not respond strongly to at least two of the three presentations of the positive control, the fly was excluded from further analysis. If projections responded to two or more presentations of the negative control, the fly was excluded from further analysis. A large ROI containing no GCaMP signal was chosen in the lateral SEZ to determine background fluorescence.

In calcium imaging experiments, the first five time points of each imaging session were discarded, leaving 40 time points for analysis with taste stimulations at time points 10, 20 and 30. The average fluorescence intensity of the background ROI was subtracted at each time point from that of the taste projection ROI. F0 was then defined as the average fluorescence intensity of the taste projection ROI post background subtraction of the first five time points. ΔF/F (%) was calculated as 100%* (F(t)-F0)/F0. Voltage imaging experiments were analyzed similarly, with ten initial time points discarded for a total of 80 time points in the analysis and tastant presentations at time points 20, 40 and 60.

### Quantification of Calcium and Voltage Imaging

Graphs were generated in GraphPad Prism. To calculate the max ΔF/F (%) of GCaMP responses, the ΔF/F(%) of the three time points centered on the peak ΔF/F (%) after the first stimulus response were averaged. The average ΔF/F (%) of the three time points immediately preceding the stimulus onset were then subtracted to account for changing baselines during imaging. Arclight data was similarly analyzed, except that five timepoints centered on the peak ΔF/F (%) and five time points prior to stimulus onset were considered. Statistical tests were performed in Prism.

### Immunohistochemistry

To visualize GRN projections with light microscopy, males of Gr64f-GAL4, Gr66a-GAL4, Ir94e-GAL4, or Ppk28-GAL4 were crossed to virgins of UAS-Syt-HA, 20XUAS-CsChrimson-mVenus (attP18). Dissection and staining were carried out by FlyLight (Gr64f-GAL4 and Gr66a-GAL4) or in house (Ir94e-GAL4 and Ppk28-GAL4) according to the FlyLight ‘IHC-Polarity Sequential Case 5’ protocol (https://www.janelia.org/project-team/flylight/protocols). Samples were imaged on an LSM710 confocal microscope (Zeiss) with a Plan-Apochromat 20×/0.8 M27 objective. Images were then registered to the 2018U template using CMTK (https://www.nitrc.org/projects/cmtk) and manually segmented with VVDViewer (https://github.com/takashi310/VVD_Viewer; Otsuna et al., 2018) in order to remove any non-specific background; Otsuna et al., 2018) in order to remove any non-specific background.

## Acknowledgements

We thank Lori Horhor, Jolie Huang, Neil Ming, and Parisa Vaziri for EM tracing contributions. This work was supported by NIH R01DC013280 (K.S.) and NIH F32DK117671 (G.S.). We thank John Bogovic for registration of EM skeletons in the 2018U template. Neuronal reconstruction for this project took place in a collaborative CATMAID environment in which 27 labs are participating to build connectomes for specific circuits. Development and administration of the FAFB tracing environment and analysis tools were funded in part by National Institutes of Health BRAIN Initiative grant 1RF1MH120679-01 to Davi Bock and Greg Jefferis, with software development effort and administrative support provided by Tom Kazimiers (Kazmos GmbH) and Eric Perlman (Yikes LLC). Peter Li, Viren Jain and colleagues at Google Research shared automatic segmentation (Li et al 2019). Members of the Scott lab and David T. Harris provided comments on the manuscript.

---

**Figure 1 - figure supplement 1.**
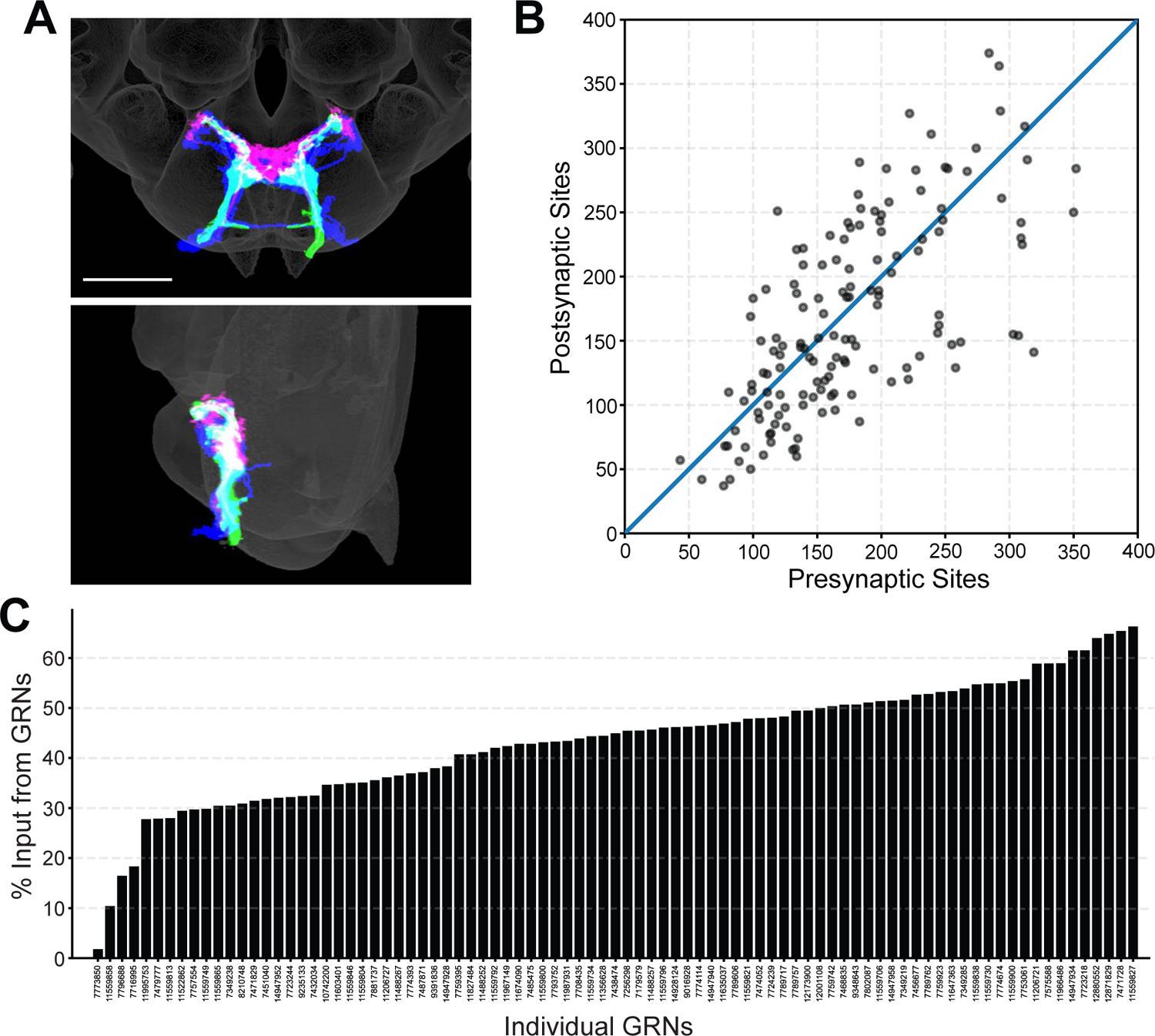
Morphology and connectivity of reconstructed GRN skeletons. (A) Overlap of reconstructed GRNs (dark blue) with the projection patterns of bitter (magenta) and sugar (green) GRNs in the 2018U template brain, frontal view (top) and sagittal view (bottom), scale bar = 50 µM. (B) Plot of pre- and post-synaptic sites for individual GRNs of the right hemisphere, denoted by grey circles. Diagonal line indicates one-to-one relationship of pre- and post-synaptic sites. (C) Percentage of GRN inputs to each GRN, for GRNs of the right hemisphere.

**Figure 2 - figure supplement 1.**
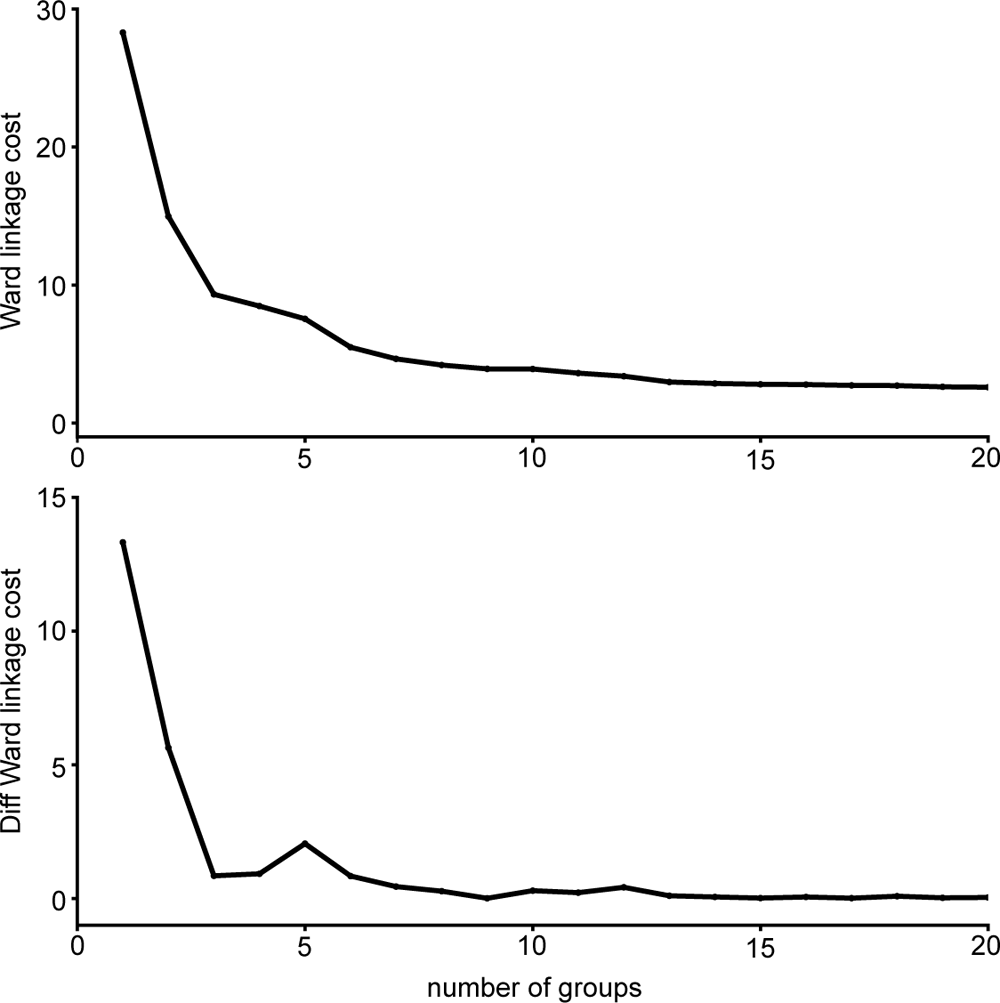
Ward’s joining cost and the differential of Ward’s joining cost for hierarchical clustering of GRNs in the right hemisphere. (top) Ward’s joining cost for clustering into groups. Ward’s joining cost declines sharply when clustering with six groups as compared to clustering with fewer than six groups. (bottom) Differential of Ward’s joining cost for clustering into groups. The differential is high when clustering into five groups or fewer but does not decline notably after six groups is reached.

**Figure 3 – figure supplement 1.**
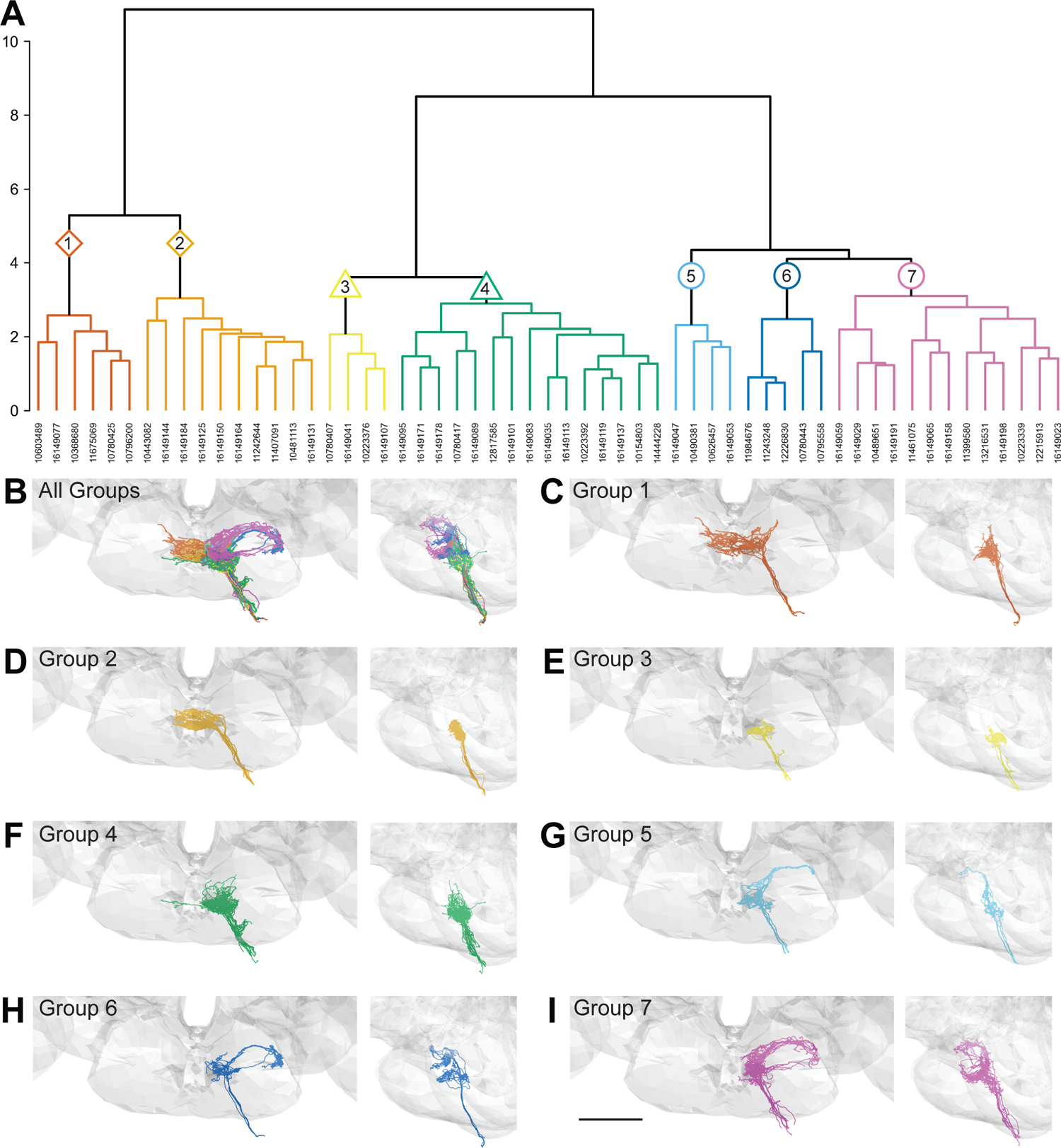
Morphology- and connectivity-based clustering generates distinct groups of GRNs. (A) Tree denoting relative similarity of GRNs based on morphology and connectivity of GRNs in the left hemisphere. (B) Frontal and sagittal view of all GRN groups, colored according to A. (C-H) Frontal and sagittal view of group 1 - group 7 GRNs, scale bar = 50 µM.

**Figure 3 - figure supplement 2.**
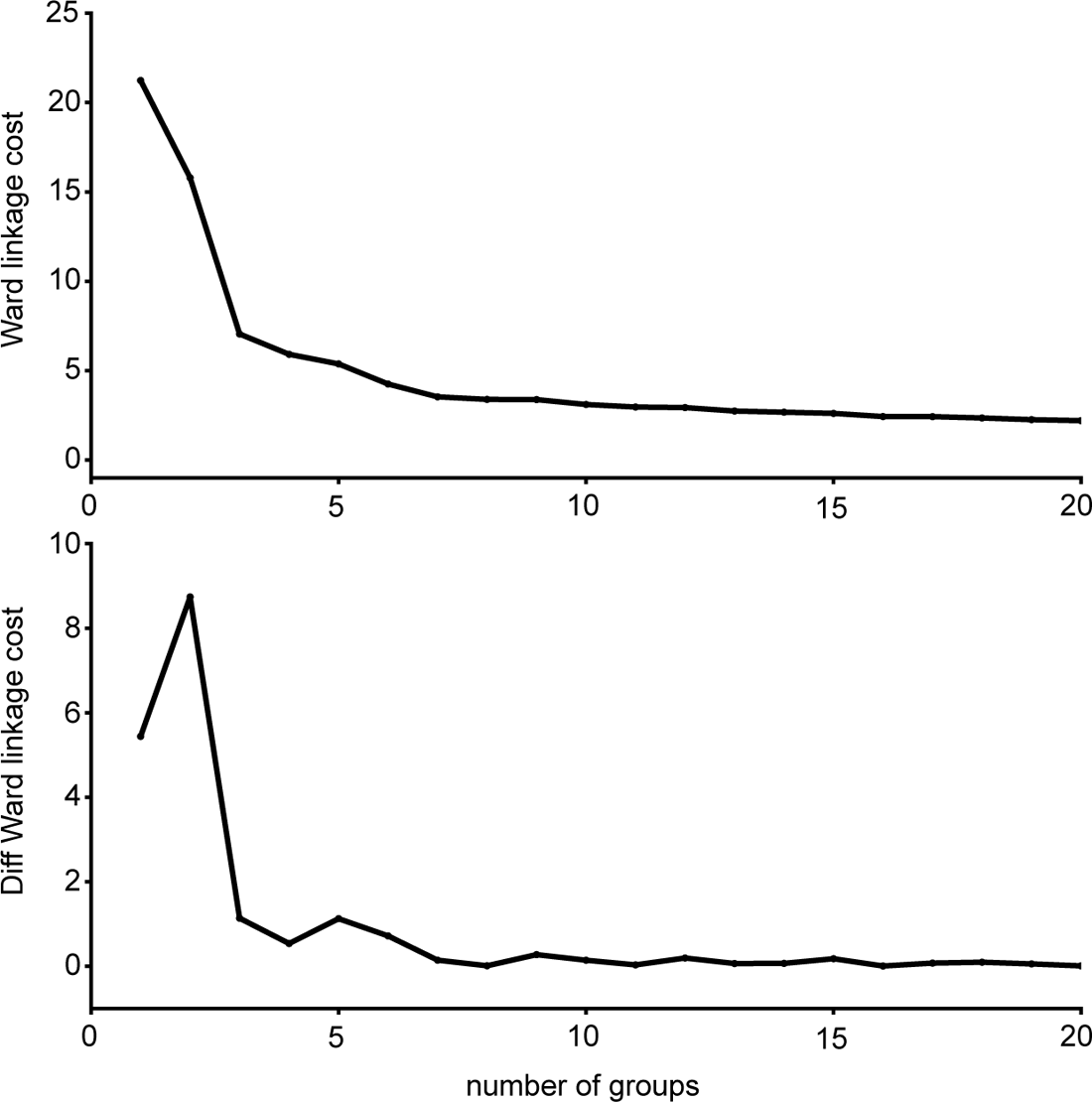
Ward’s joining cost and the differential of Ward’s joining cost for hierarchical clustering of GRNs in the left hemisphere. (top) Ward’s joining cost for clustering into groups. Ward’s joining cost declines sharply when clustering with seven groups as compared to clustering with fewer than seven groups. (bottom) Differential of Ward’s joining cost for clustering into groups. The differential is high when clustering into six groups or fewer but does not decline notably after seven groups is reached.

**Figure 5 - figure supplement 1.**
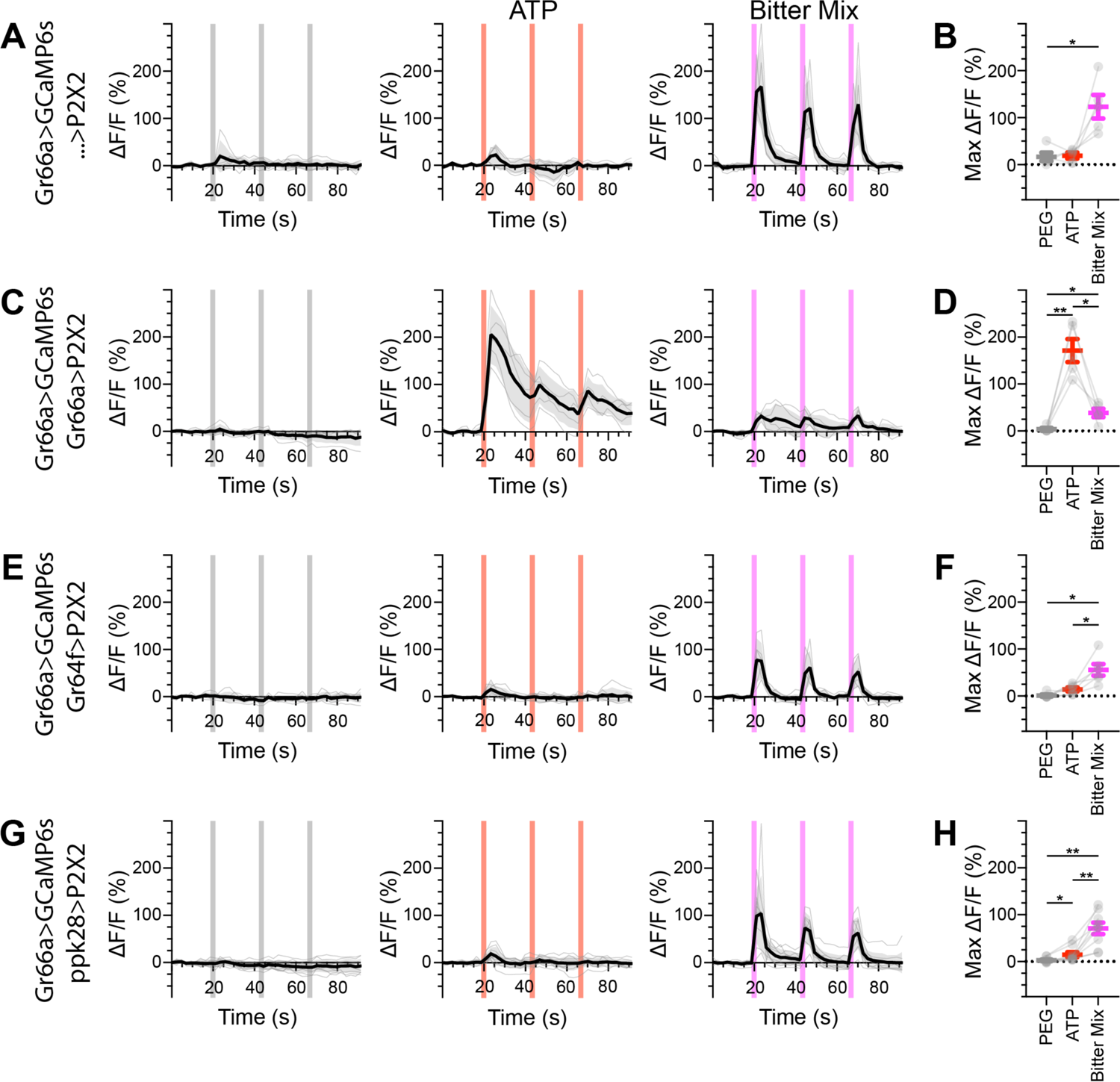
Bitter GRNs do not respond to the activation of other GRN classes in fed flies. (A, B) Calcium responses of bitter GRNs expressing GCaMP6s in a UAS-P2X2 background to proboscis presentation of PEG as a negative control, ATP, or a mixture of denatonium and caffeine, which are bitter compounds, as a positive control, GCaMP6s ΔF/F traces (A) and maximum ΔF/F graph (B), n = 5. (C, D) Calcium responses of bitter GRNs expressing GCaMP6s and P2X2 to PEG, ATP, or bitter delivery, ΔF/F traces (C) and maximum ΔF/F graph (D), n = 5. (E, F) GCaMP6s responses of bitter GRNs in flies expressing P2X2 in sugar GRNs to PEG, ATP, and bitter, ΔF/F traces (E) and maximum ΔF/F graph (F), n = 6. (G, H) GCaMP6s responses of bitter GRNs in flies expressing P2X2 in water GRNs to delivery of PEG, ATP, or bitter to the proboscis, ΔF/F traces (G) and maximum ΔF/F graph (H), n = 9. Period of stimulus presentation is indicated by shaded bars, 3 stimulations/fly. Traces of individual flies are shown in grey, the average in black, with the SEM indicated by the grey shaded area. Repeated measures ANOVA with Tukey’s multiple comparisons test, *p<0.05, **p<0.01.

**Figure 5 - figure supplement 2.**
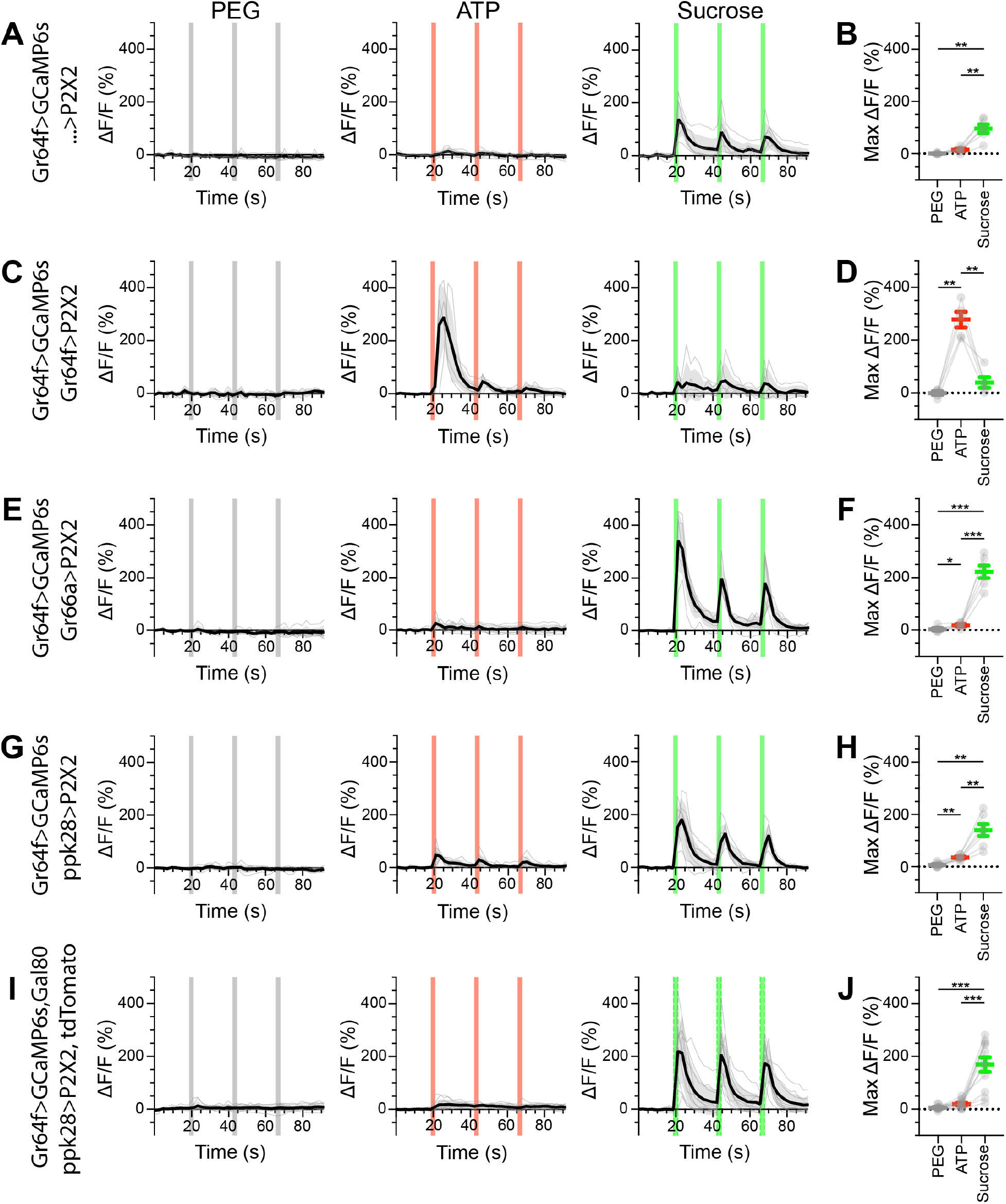
Sugar GRNs do not respond to the activation of other GRN classes in fed flies. (A, B) Calcium responses of sugar GRNs expressing GCaMP6s in a UAS-P2X2 background to proboscis presentation of PEG as a negative control, ATP, or sucrose as a positive control, GCaMP6s ΔF/F traces (A) and maximum ΔF/F graph (B), n = 6. (C, D) Calcium responses of sugar GRNs expressing GCaMP6s and P2X2 to PEG, ATP, or sucrose delivery, ΔF/F traces (C) and maximum ΔF/F graph (D), n = 5. (E, F) GCaMP6s responses of sugar GRNs in flies expressing P2X2 in bitter GRNs to PEG, ATP, and sucrose, ΔF/F traces (E) and maximum ΔF/F graph (F), n = 6. (G, H) GCaMP6s responses of sugar GRNs in flies expressing P2X2 in water GRNs to PEG, ATP, or sucrose presentation, ΔF/F traces (G) and maximum ΔF/F graph (H), n = 7. (I, J) GCaMP6s responses of sugar GRNs in flies expressing P2X2 in water GRNs and Gal80 in sugar GRNs to inhibit P2X2 misexpression to PEG, ATP, or sucrose presentation, ΔF/F traces (I) and maximum ΔF/F plots (J), n = 11. Period of stimulus presentation is indicated by shaded bars, 3 stimulations/fly. Data from first stimulation of C and K is shown in Figure 4A-D. Traces of individual flies are shown in grey, the average in black, with the SEM indicated by the grey shaded area. Repeated measures ANOVA with Tukey’s multiple comparisons test *p<0.05, **p<0.01, ***p<0.001.

**Figure 5 - figure supplement 3.**
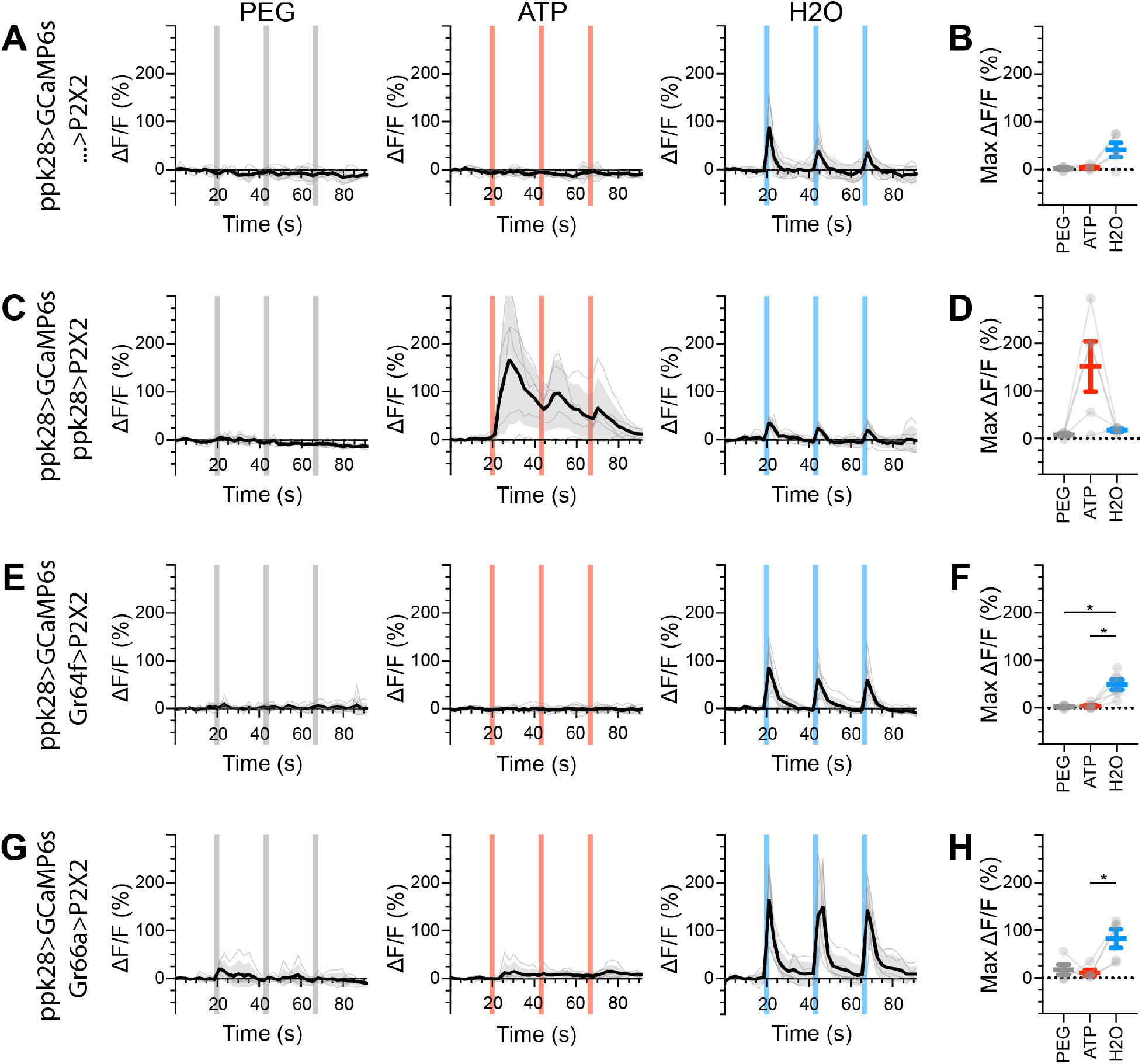
Water GRNs do not respond to the activation of other GRN classes in fed flies. (A, B) Calcium responses of water GRNs expressing GCaMP6s in a UAS-P2X2 background to proboscis presentation of PEG as a negative control, ATP, or water as a positive control, GCaMP6s ΔF/F traces (A) and maximum DF/F graph (B), n = 5. (C, D) Calcium responses of water GRNs expressing GCaMP6s and P2X2 to PEG, ATP, or water delivery, ΔF/F traces (C) and maximum ΔF/F graph (D), n = 5. (E, F) GCaMP6s responses of water GRNs in flies expressing P2X2 in sugar GRNs to PEG, ATP, and water, DF/F traces (E) and maximum DF/F graph (F), n = 6. (G, H) GCaMP6s responses of water GRNs in flies expressing P2X2 in bitter GRNs upon PEG, ATP, or water presentation, ΔF/F traces (G) and maximum ΔF/F graph (H), n = 5. Period of stimulus presentation is indicated by shaded bars, 3 stimulations/fly. The first response in C and E is shown in Figure 4G-J. Traces of individual flies are shown in grey, the average in black, with the SEM indicated by the grey shaded area. Repeated measures ANOVA with Tukey’s multiple comparisons test *p<0.05.

**Figure 5 - figure supplement 4.**
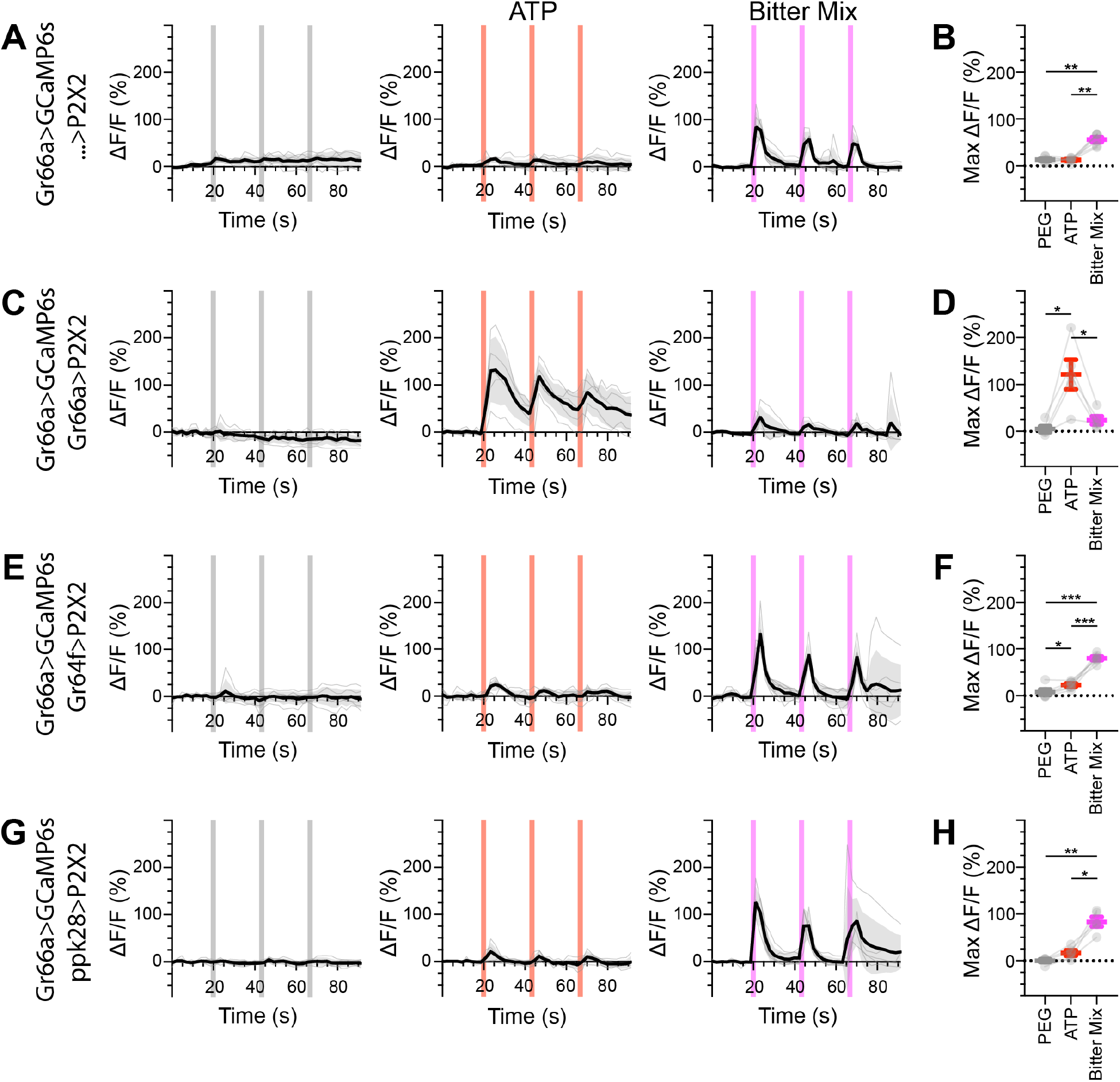
Bitter GRNs do not respond to the activation of other GRN classes in food-deprived flies. (A, B) Calcium responses of bitter GRNs expressing GCaMP6s in a UAS-P2X2 background to proboscis presentation of PEG as a negative control, ATP, or a mixture of the bitter compounds denatonium and caffeine as a positive control, GCaMP6s ΔF/F traces (A) and maximum ΔF/F graph (B), n = 6. (C, D) Calcium responses of bitter GRNs expressing GCaMP6s and P2X2 to PEG, ATP, or bitter delivery, ΔF/F traces (C) and maximum ΔF/F graph (D), n = 5. (E, F) GCaMP6s responses of bitter GRNs in flies expressing P2X2 in sugar GRNs to PEG, ATP, and bitter, ΔF/F traces (E) and maximum ΔF/F graph (F), n = 6. (G, H) GCaMP6s responses of bitter GRNs in flies expressing P2X2 in water GRNs to delivery of PEG, ATP, or bitter, ΔF/F traces (G) and maximum ΔF/F graph (H), n = 5. Period of stimulus presentation is indicated by shaded bars, 3 stimulations/fly. Flies were food-deprived for 23-26 hours. Traces of individual flies are shown in grey, the average in black, with the SEM indicated by the grey shaded area. Repeated measures ANOVA with Tukey’s multiple comparisons test, *p<0.05, **p<0.01, ***p<0.001.

**Figure 5 - figure supplement 5.**
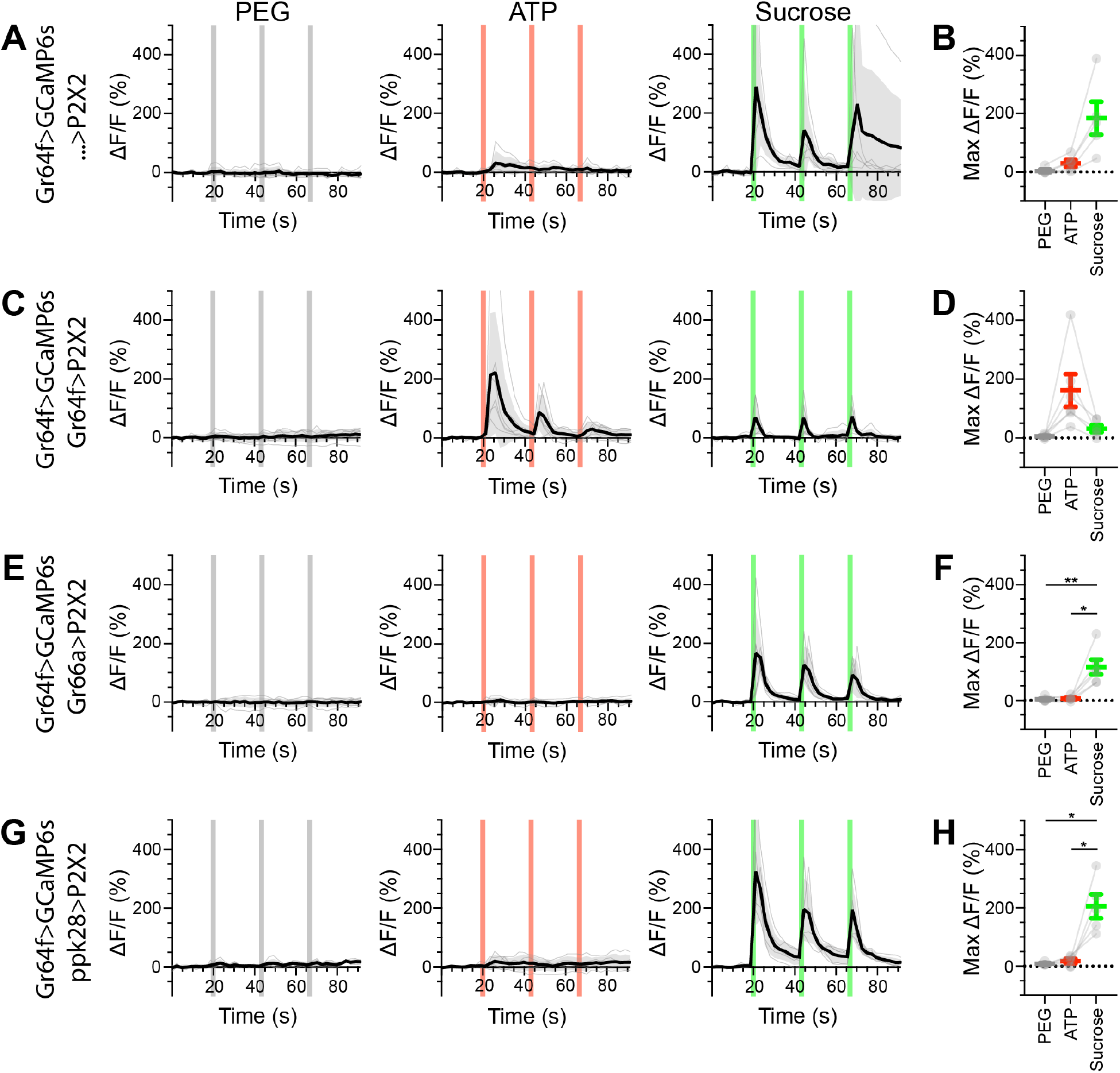
Sugar GRNs do not respond to the activation of other GRN classes in food-deprived flies. (A, B) Calcium responses of sugar GRNs expressing GCaMP6s in a UAS-P2X2 background to proboscis presentation of PEG as a negative control, ATP, or sucrose as a positive control, GCaMP6s ΔF/F traces (A) and maximum ΔF/F graph (B), n = 5. (C, D) Calcium responses of sugar GRNs expressing GCaMP6s and P2X2 to PEG, ATP, or sucrose delivery, ΔF/F traces (C) and maximum ΔF/F graph (D), n = 6. (E, F) GCaMP6s responses of sugar GRNs in flies expressing P2X2 in bitter GRNs to PEG, ATP, and sucrose, ΔF/F traces (E) and maximum ΔF/F graph (F), n = 6. (G, H) GCaMP6s responses of sugar GRNs in flies expressing P2X2 in water GRNs to PEG, ATP, and sucrose presentation to the proboscis, ΔF/F traces (G) and maximum ΔF/F graph (H), n = 5. Period of stimulus presentation is indicated by shaded bars, 3 stimulations/fly. Flies were food-deprived for 23-26 hours. Traces of individual flies are shown in grey, the average in black, with the SEM indicated by the grey shaded area. Repeated measures ANOVA with Tukey’s multiple comparisons test *p<0.05, **p<0.01.

**Figure 5 - figure supplement 6.**
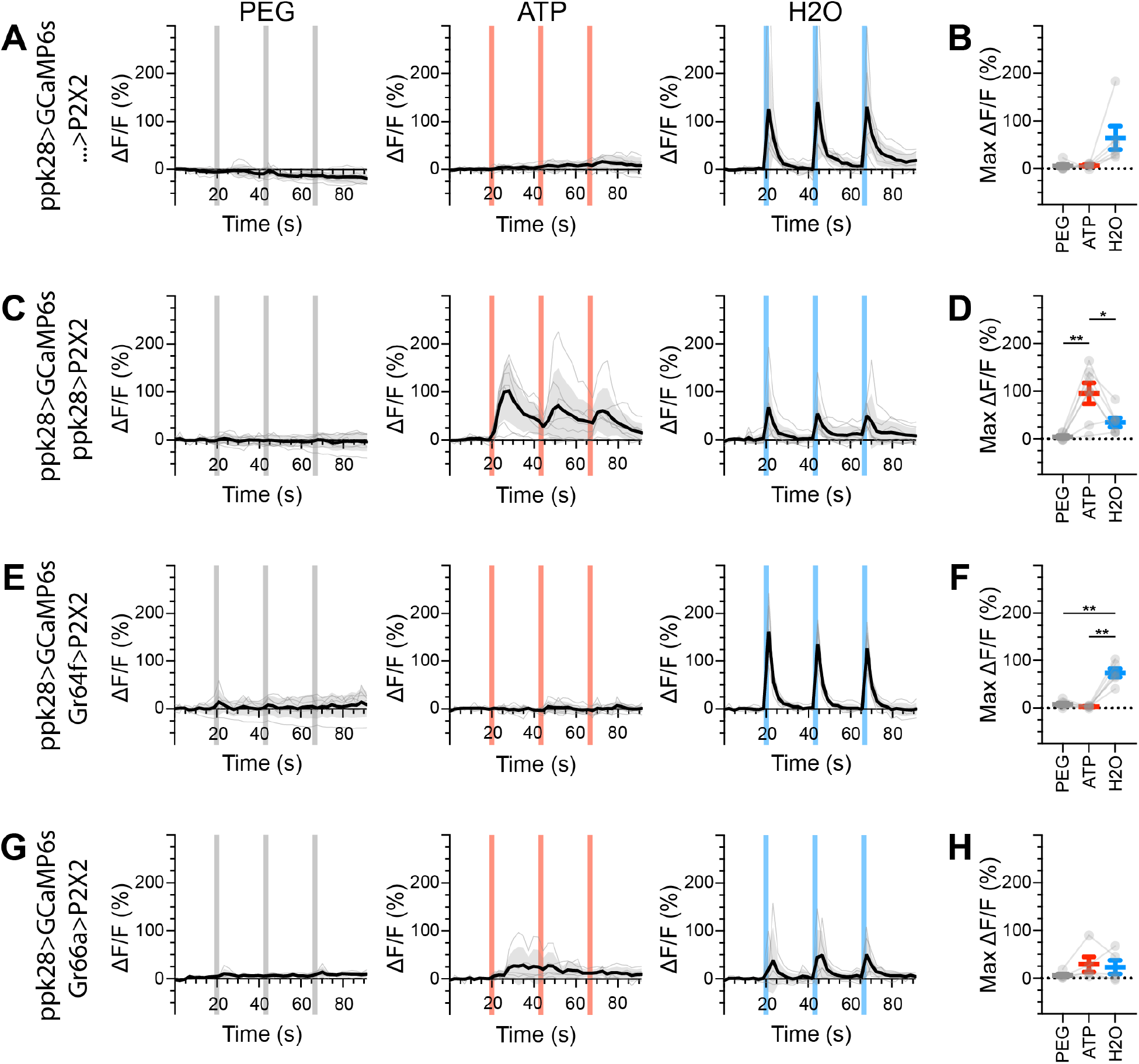
Water GRNs do not respond to the activation of other GRN classes in food-deprived flies. (A, B) Calcium responses of water GRNs expressing GCaMP6s in a UAS-P2X2 background to proboscis presentation of PEG as a negative control, ATP, or water as a positive control, GCaMP6s ΔF/F traces (A) and maximum ΔF/F graph (B), n = 6. (C, D) Calcium responses of water GRNs expressing GCaMP6s and P2X2 to PEG, ATP, or water delivery, ΔF/F traces (C) and maximum ΔF/F graph (D), n = 7. (E, F) GCaMP6s responses of water GRNs in flies expressing P2X2 in sugar GRNs to PEG, ATP, and water, ΔF/F traces (E) and maximum ΔF/F graph (F), n = 6. (G, H) GCaMP6s responses of water GRNs in flies expressing P2X2 in bitter GRNs to PEG, ATP, and water delivery, ΔF/F traces (G) and maximum ΔF/F graph (H), n = 5. Period of stimulus presentation is indicated by shaded bars, 3 stimulations/fly. Flies were food-deprived for 23-26 hours. Traces of individual flies are shown in grey, the average in black, with the SEM indicated by the grey shaded area. Repeated measures ANOVA with Tukey’s multiple comparisons test *p<0.05, **p<0.01.

**Figure 5 - figure supplement 7.**
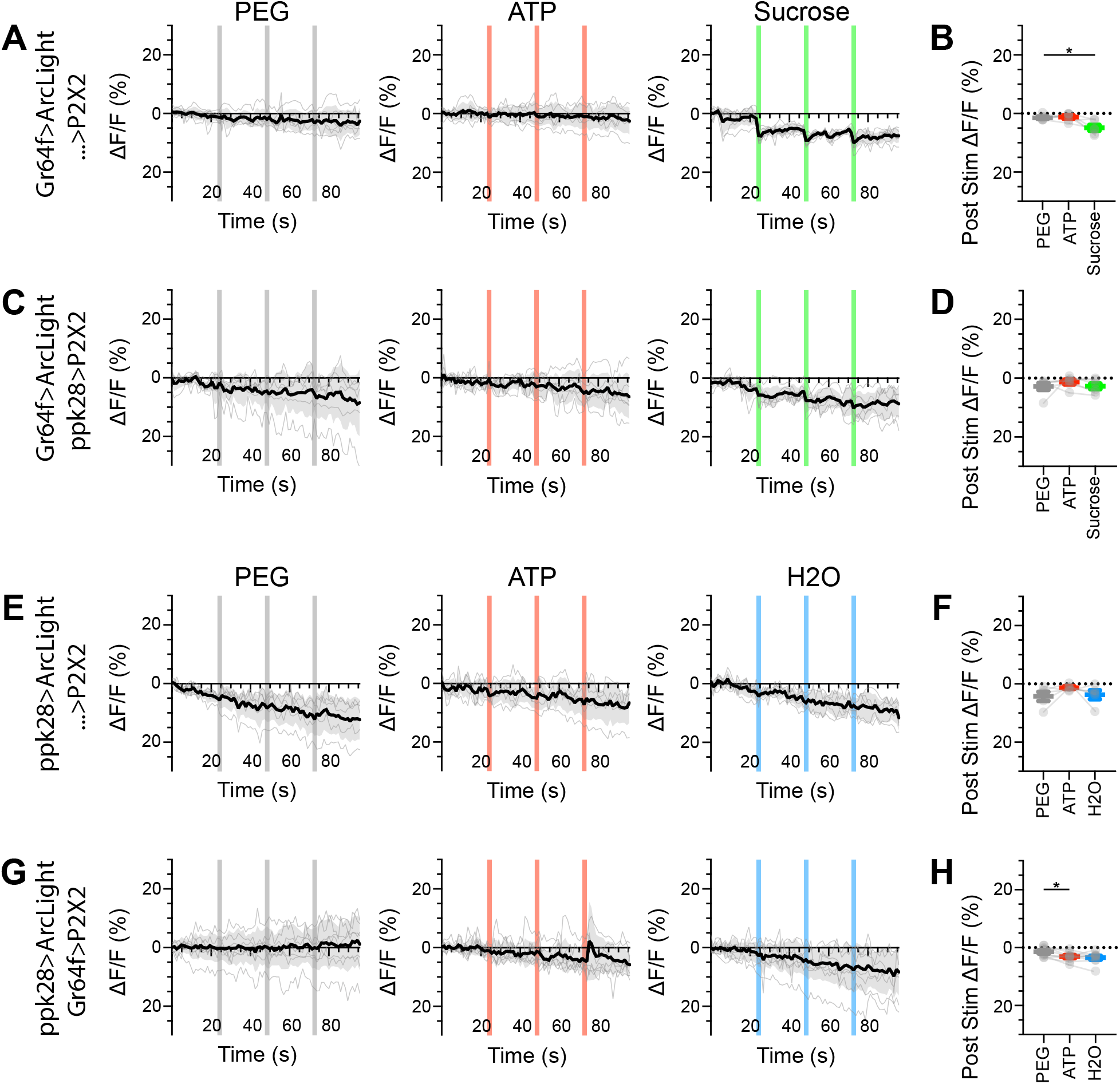
Sugar and water GRNs do not show voltage responses upon reciprocal activation. (A, B) ArcLight responses of sugar GRNs in a UAS-P2X2 background to proboscis presentation of PEG as a negative control, ATP, or sucrose as a positive control. ArcLight fluorescence traces (ΔF/F) (A) and maximum ΔF/F post stimulus presentation (B), n = 6. (C, D) ArcLight responses of sugar GRNs in flies expressing P2X2 in water GRNs to PEG, ATP, and sucrose delivery, ΔF/F traces (C) and maximum ΔF/F graph (D), n = 6. (E, F) ArcLight responses of water GRNs in a UAS-P2X2 background to proboscis delivery of PEG, ATP, and water (positive control), ΔF/F traces (E) and maximum ΔF/F graph (F), n = 5. (G, H) ArcLight responses of water GRNs in flies expressing P2X2 in sugar GRNs to PEG, ATP, and water delivery, ΔF/F traces (G) and maximum ΔF/F graph (H), n = 9. Period of stimulus presentation is indicated by shaded bars, 3 stimulations/fly. The first response in C and G is shown in Figure 2-4 E, F, K, L. Traces of individual flies to three taste stimulations are shown in grey, the average in black, with the SEM indicated by the grey shaded area. Repeated measures ANOVA with Tukey’s multiple comparisons test, *p<0.05.

